# FB5P-seq-mAbs: monoclonal antibody production from FB5P-seq libraries for integrative single-cell analysis of B cells

**DOI:** 10.1101/2024.10.01.616021

**Authors:** Sakina Ado, Chuang Dong, Noudjoud Attaf, Myriam Moussa, Agathe Carrier, Pierre Milpied, Jean-Marc Navarro

## Abstract

Parallel analysis of phenotype, transcriptome and antigen receptor sequence in single B cells is a useful method for tracking B cell activation and maturation during immune responses. However, in most cases, the specificity and affinity of the B cell antigen receptor cannot be inferred from its sequence. Antibody cloning and expression from single B cells is then required for functional assays. Here we propose a method that integrates FACS-based 5’-end single-cell RNA sequencing (FB5P-seq) and monoclonal antibody cloning for integrative analysis of single B cells. Starting from a cell suspension, single B cells are FACS-sorted into 96-well plates for reverse transcription, cDNA barcoding and amplification. A fraction of the single-cell cDNA is used for preparing 5’-end RNA-seq libraries that are sequenced for retrieving transcriptome-wide gene expression and paired BCR sequences. The archived cDNA of selected cells of interest is used as input for cloning heavy and light chain variable regions into antibody expression plasmid vectors. The corresponding monoclonal antibodies are produced by transient transfection of a eukaryotic producing cell line and purified for functional assays. We provide detailed step-by-step instructions and describe results obtained on ovalbumin-specific murine germinal center B cells after immunization. Our method is robust, flexible, cost-effective, and applicable to different B cell types and species. We anticipate it will be useful for mapping antigen specificity and affinity of rare B cell subsets characterized by defined gene expression and/or antigen receptor sequence.

## INTRODUCTION

In the immune response to pathogens and vaccines, antigen-responsive B cells undergo series of cellular and molecular maturation events that are required for long term immune protection and memory (1). On the cellular side, activated B cells divide, migrate, and evolve through distinct intermediate differentiation stages that ultimately give rise to long-lived antibody-producing plasma cells and memory B cells (2). On the molecular side, within each responding B cell, the genetic *loci* encoding B cell receptor (BCR) immunoglobulin heavy (IGH) and light (IGK/L) chains may undergo class switch recombination (CSR) (3) and somatic hypermutation (SHM) (4), providing opportunities for improving the function and affinity of antigen-specific antibodies produced throughout the current and future immune responses (5). In those parallel cellular and molecular evolution processes, every antigen-responsive B cell may generate a diverse progeny of clonally related daughter B cells expressing unique combinations of functional properties and BCR affinities. Thus, methods that enable integrative analysis of B cell phenotype, transcriptome, BCR sequence, and BCR affinity at the single-cell level, are useful tools when studying B cell immune responses (6).

In the past few years, two types of single-cell RNA sequencing (scRNA-seq) techniques have been applied to B cells for integrative analysis of transcriptome and BCR sequence: plate-based scRNA-seq of B cells sorted by flow cytometry, either with full-length (Smart-seq2) (7,8) or 5’-end sequencing (FB5P-seq) (9); and droplet-based 10x Genomics 5’-end scRNA-seq (10). Both techniques may link B cell transcriptome and BCR sequence with antigen-specificity, provided B cells are incubated with labeled antigen (fluorescent antigen for FACS-based methods (11), DNA-barcoded antigen for droplet-based method (12)) before analysis. However, those approaches are not suitable for B cells expressing low amounts of surface BCR (e.g. plasma cells), and preclude extensive analyses of BCR specificity, affinity and function.

Antibody cloning and production from single FACS-sorted memory B cells or plasmablasts (e.g. protocols described in (13–15)), has been the method of choice for discovering and characterizing naturally occurring antigen-specific antibodies from infected individuals, notably in the fields of HIV (16) and SARS-CoV2 (17) research. Such methods, applied to animal models of protein vaccination or infection, have also contributed to our basic understanding of the role of BCR affinity during germinal center (GC) B cell responses (18,19). However, in those studies, the cellular characteristics of B cells from which monoclonal antibodies (mAbs) are cloned can only be inferred from the expression of a few surface markers or of fluorescent reporters.

Here we describe FB5P-seq-mAbs, a method that bridges plate-based 5’-end scRNA-seq (FB5P-seq) with recombinant monoclonal antibody cloning and production (**Figure 1**), and illustrate its use for characterizing antigen-responding GC B cells after chicken ovalbumin (OVA) immunization in mice.

**Figure 1.**
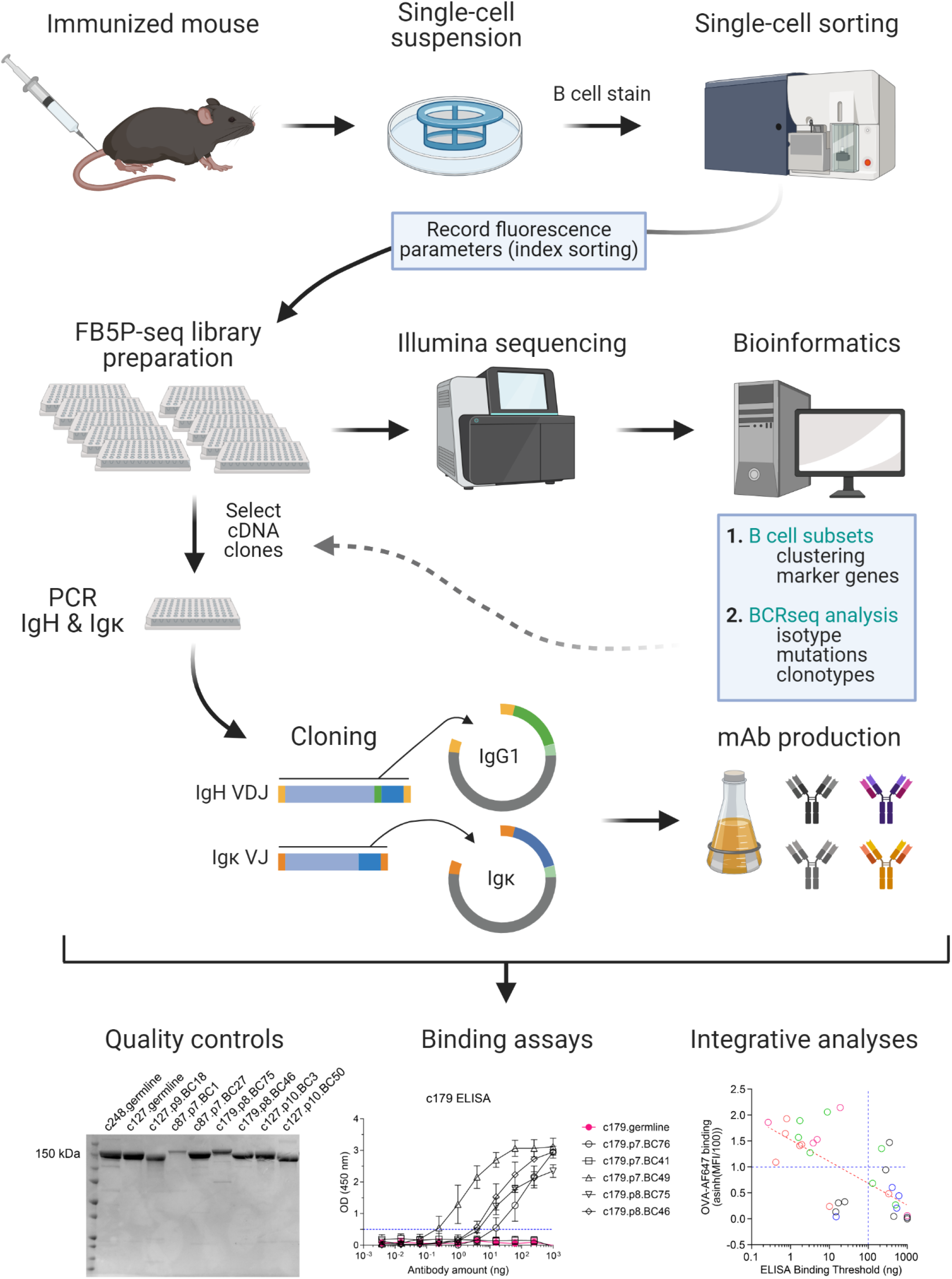
Workflow of FB5P-seq-mAbs. Starting from an immunized mouse, single B cells of interest are sorted by FACS for 5’-end single-cell RNA-seq library preparation with the FB5P-seq protocol (9), recording fluorescence intensity parameters of each sorted cell by index sorting. FB5P-seq libraries from multiple 96-well plates are pooled, sequenced, and analyzed to identify B cell subsets, recover paired BCR heavy and light chain sequences, and integrate both information types. For selected cells of interest, the remaining single-cell amplified cDNA in archived 96-well PCR plates is used as starting material for PCR amplification of IgH VDJ and Igκ VJ sequences, and cloning into mouse IgG1 and Igκ expression vectors, respectively. IgG1 and Igκ expression plasmids are co-transfected into a eukaryotic cell line for production of recombinant mAb in the culture supernatant. After purification, recombinant IgG1/κ mAbs are available for functional assays and integrative analyses. Created with BioRender.com.

## MATERIALS AND EQUIPMENT

### Antibodies

| Reagent or resource | Source | Identifier |
| --- | --- | --- |
| CD138-BV711 | Biolegend | cat.no.142519 |
| CD19-PE-Dazzle594 | Biolegend | cat.no. 115553 |
| CD38-PE | Biolegend | cat.no. 356608 |
| CD3-APC-Cy7 | Biolegend | cat.no.100222 |
| CXCR4-PerCP-eF710 | ThermoFisher Scientific | cat.no. 46-9991-82 |
| GL7-BV421 | Biolegend | cat.no.144614 |
| Gr1-APC-Cy7 | Biolegend | cat.no.108424 |
| Mouse Fc block | Biolegend | cat.no.101320 |
| OVA-AF 647 | ThermoFisher Scientific | cat.no. O34784 |
| HRP-coupled anti-mouse IgG secondary antibody | ThermoFisher Scientific | cat.no. A24512 |

### Cell lines, mouse strains, bacteria

| Reagent or resource | Source | Identifier |
| --- | --- | --- |
| Aicda-Cre-ERT2 x Rosa26-lox-STOP-lox-eYFP | Le Gallou et al. (20) |  |
| E.Coli JM109 | Promega | cat.no. L2005 |
| Expi293TM | ThermoFisher Scientific | cat.no. A14527 |

### Chemicals, peptides and recombinant proteins

| Reagent or resource | Source | Identifier |
| --- | --- | --- |
| 1Kb plus DNA ladder | ThermoFisher Scientific | cat.no. 10787018 |
| 2N sulfuric acid | ThermoFisher Scientific | cat.no. N600 |
| 2-Propanol | Sigma-Aldrich | cat.no. 109634 |
| Acetic acid | VWR | cat.no. 20104.334 |
| Agarose | Life Technologies | cat.no. 16500500 |
| Agel-HF | New England Biolabs | cat.no. R3552S |
| Alkaline Phosphatase | New England Biolabs | cat.no. M0525S |
| Alum | ThermoFisher Scientific | cat.no. 77161 |
| Ampicillin | Sigma-Aldrich | cat.no. A9518 |
| Arachidic acid oil | Sigma-Aldrich | cat.no. A3631 |
| Betaine | Sigma-Aldrich | cat.no. B0300 |
| Boric Acid | Carlo Erba | cat.no. 600641 |
| Bromophenol Blue | Sigma-Aldrich | Cat no. B0126-25G |
| Bovine Serum Albumin | Sigma-Aldrich | cat.no. 10735078001 |
| BsiwI | New England Biolabs | cat.no. R0553L |
| Clean NGS beads | Proteigene | cat.no. CNGS-0050 |
| CutSmart Buffer | New England Biolabs | cat.no. B7203S |
| Dimethyl sulfoxide | Merck | cat.no. 276855-100ML |
| dNTP | ThermoFisher Scientific | cat.no. 10297018 |
| EDTA | ThermoFisher Scientific | cat.no. 15575020 |
| Ethanol | Sigma-Aldrich | cat.no. M3148-100ML |
| Ethidium bromide | Merck | cas.no. E1510 |
| Expi293™ medium | ThermoFisher Scientific | cat.no. A1435101 |
| Fetal Calf Serum | Gibco | cat.no. 10270-106 |
| Glucose | Merck | cat.no. 49159-1KG |
| Glycine | ThermoFisher Scientific | cat no. 15527-013 |
| GoTaq G2 Flexi enzyme | Promega | cat no. M7801 |
| HCl | Dutscher | cat no. 524526 |
| KAPA HiFi ReadyMix | Roche Diagnostics | cat.no. 7958935001 |
| LB Agar | ThermoFisher Scientific | cat.no. 22700025 |
| LB medium | ThermoFisher Scientific | cat.no. 12780052 |
| Live/Dead | ThermoFisher Scientific | cat.no. L34957 |
| Protein ladder | Biorad | cat.no. 1610373 |
| Methanol | Dutscher | cat.no. 412383 |
| MgCl<sub>2</sub> | Sigma-Aldrich | cat.no. M1028 |
| NaCl | Sigma-Aldrich | cat.no. S6546-1L |
| NEBuffer 2 | New England Biolabs | cat.no. B7202 |
| NEBuffer 3.1 | New England Biolabs | cat.no. B7203S |
| Nupage novex 4-12% BT midi gel | ThermoFisher Scientific | cat.no. WG1401BOX |
| NuPAGE™ MOPS SDS Running Buffer (20X) | Life tech | cat.no. NP0001 |
| OVA antigen | Invivogene | cat.no. vac-stova |
| OVA stock | ThermoFisher Scientific | cat.no. 77120 |
| PBS 10 X | Life Technologies | cat.no. 14200067 |
| PCR-grade H<sub>2</sub>O | Qiagen | cat.no. 129114 |
| Penicillin-streptomycin | ThermoFisher Scientific | cat.no. 15140122 |
| Polyethylene glycol 3350 | Merck | cat.no. P4338-500G |
| Polyethyleneimine | Sigma-Aldrich | cat.no. 408727-100ML |
| Protein G beads | Sigma-Aldrich | cat.no. GE17-0618-01 |
| Rnase OUT | ThermoFisher Scientific | cat.no. 10777019 |
| Laemmli SDS-Sample Buffer 4X, Non-Reducing | Cliniscience | cat.no. 10570018-1 |
| Sample Buffer Laemmli 2x Concentrate | Sigma-Aldrich | cat.no. S3401-10VL |
| SOC medium | Invitrogen | cat.no. 15544-034 |
| Sodium azide | Merck | cat.no. S2002-5G |
| SuperScript II | ThermoFisher Scientific | cat.no. 18064014 |
| T4 DNA polymerase | New England Biolabs | cat.no. M0203S |
| Tamoxifen | Sigma-Aldrich | cat.no. T5648-1G |
| TMB Solution | ThermoFisher Scientific | cat.no. 34021 |
| Tris base | Life Technologies | cat.no. 15504020 |
| Triton X-100 | Merck | cat.no. 93443-100ML |
| Trypan blue | Merck | cat.no. T8154-20ML |
| Tryptone | Merck | cas.no. T7293-250G |
| Tween 20 | Merck | cat.no. P9416-100ML |
| Valproic acid | Merck | cat.no. V0033000 |
| Xhol | New England Biolabs | cat.no. R0146L |
| Yeast extract | Merck | cat.no. Y1625-250G |

### Oligonucleotides

| Reagent or resource | Source | Identifier |
| --- | --- | --- |
| (dT)30_Smarter (see table 1) | IDT | *Attaf et al* (9) |
| ERCC spike-in Mix (see table 1) | ThermoFisher Scientific | cat.no. 4456740 |
| i5 primer S5xx | Illumina | cat.no. FC-131-2001 |
| i7 primers mix | IDT | *Attaf et al* (9) |
| i7_BCx primer | IDT | *Attaf et al* (9) |
| Multiplex Forward Primers | IDT | *Supplementary Table 1* |
| Multiplex Reverse Primers | IDT | *Supplementary Table 1* |
| PCR_Satija (see table 1) | IDT | *Attaf et al (9)* |
| Screening Forward Primers (see table 4) | IDT | *Supplementary Table 2* |
| Screening Reverse Primers (see table 4) | IDT | *Supplementary Table 2* |
| SmarterR (see table 1) | IDT | *Attaf et al (9)* |
| TSO_BCx_UMI5_TATA | IDT | *Attaf et al (9)* |

### Commercial assays

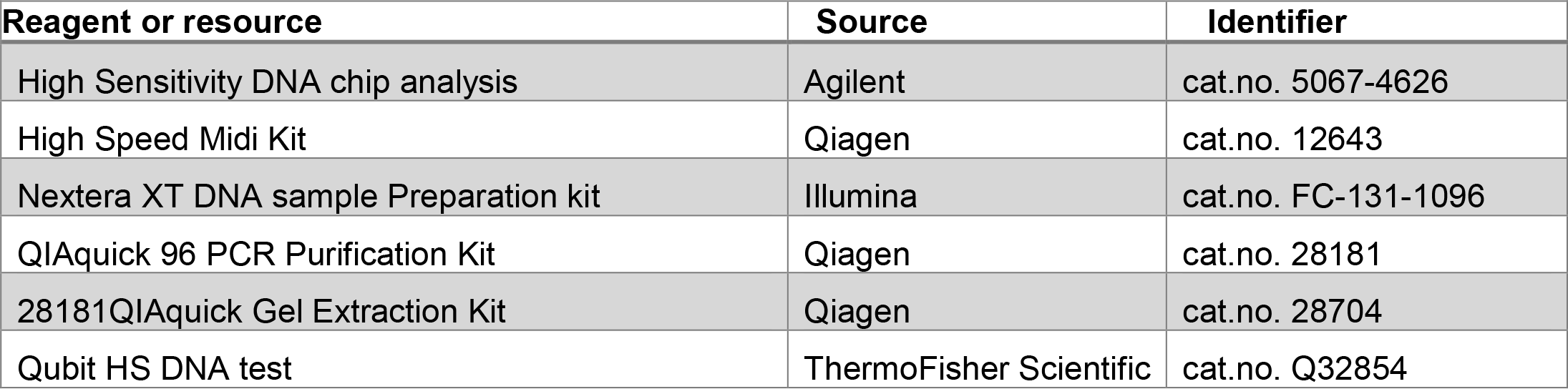

### Equipment

| **Reagent or resource** | **Source** | **Identifier** |
| --- | --- | --- |
| 8-channel micropipette 10 µl | Starlab | cat.no. S7108-0510 |
| 8-channel micropipette 100 µl | Starlab | cat.no. S7108-1100 |
| 8-channel micropipette 300 µl | Starlab | cat.no. S7108-3300 |
| BD Influx™ Cell Sorter | BD Biosciences | NA |
| Benchtop plate centrifuge | Starlab | cat.no. N2631-0008 |
| Incubator | Fisher Scientific | cat.no. 12815883 |
| DynaMag™-96 | ThermoFisher Scientific | cat.no. 12331D |
| Nanodrop | ThermoFisher Scientific | cat.no. ND-ONE |
| p10 micropipette | Starlab | cat.no. S7100-0510 |
| p1000 micropipette | Starlab | cat.no. S7110-1000 |
| p200 micropipette | Starlab | cat.no. S7100-2200 |
| Plate reader | BMG LABTECH | NA |
| Shaker | VWR | cat.no. 444-4227 |
| Sonicator | Kinematica | cat.no. HS1200 E |
| Thermal Cycler | ThermoFisher Scientific | cat.no. A24811 |
| UV light | Dutscher | cat.no. 5207326 |
| Vortex | Dutscher | cat.no. 79008 |

### Consumables

| Reagent or resource | Source | Identifier |
| --- | --- | --- |
| 0.22-micron steritop filters | Millipore | cat.no. SCGPT01RE |
| 0.3 ml syringe | Dutscher | cat.no. 324826 |
| 1.5 ml tube | Eppendorf | cat.no. 30108051 |
| 10 ml serological pipette | Becton dickinson | cat.no. 357551 |
| 10 µl filter tips | Starlab | cat.no. S1120-3810 |
| 1000 µl filter tips | Dutscher | cat.no. 134000CL |
| 15 ml tube | Dutscher | cat.no. 352096 |
| 200 µl filter tips | Sarstedt | cat.no. 3070279 |
| 25 ml serological pipette | Sarstedt | cat.no. 86.1685.001 |
| 30 KD amicon | Merck | cat.no. UFC503024 |
| 5 ml FACS tubes | Dutscher | cat.no. 352054 |
| 5 ml serological pipette | Sarstedt | cat.no. 86.1253.001 |
| 50 ml tube | Sarstedt | cat.no. 62.547.254 |
| 70 µm cell strainer | Sarstedt | cat.no. 83.3945.070 |
| 8-tube PCR strip | Merck | cat.no. EP0030124359 |
| 96 round bottom | Dutscher | cat.no. 353077 |
| 96-well ELISA plates | ThermoFisher Scientific | cat.no. 467320 |
| 96-well PCR plates | ThermoFisher Scientific | cat.no. 4483485 |
| Adhesive film | Life Technologies | cat.no. 4306311 |
| Aluminum foil | VWR | cat.no. 291-0045 |
| Chromatography columns | Sigma | cat.no. 7311550 |
| Dissection scissors | Dutscher | cat.no. HWB 002-11 |
| Flasks with vented caps | Dutscher | cat.no. 355121B |
| Gavage syringe | Fine Science Tools | cat.no. 18061-22 |
| MicroAmp EnduraPlate Optical 96-Well plates | ThermoFisher Scientific | cat.no. 4483485 |
| Parafilm | Merck | cat.no. 291-1213 |
| PCR tube | Eppendorf | cat.no. 30124359 |
| Petri Dish | Dutscher | cat.no. 633180 |
| Pre-Separation Filter | Miltenyi Biotec | cat.no. 130-041-407 |
| Reservoir for multichannel pipettes | Sigma | cat.no. 4870 |
| Stericup | Millipore | cat.no. S2GPU05RE |
| Tube DNA LoBind 0.5 ml | Merck | cat.no. EP0030108035 |
| Tube DNA LoBind 1.5 ml | Merck | cat.no. EP0030108051 |

### Software

| Reagent or resource | Source | Identifier |
| --- | --- | --- |
| Adobe Illustrator CC 2019 | Adobe | https://www.adobe.com/fr/products/illustrator.html |
| Blastn | NCBI | https://blast.ncbi.nlm.nih.gov/ |
| BD FACS sort | BD Biosciences | NA |
| FlowJo v10.8 | BD Biosciences | https://www.flowjo.com/ |
| ggplot2 | NA | https://ggplot2.tidyverse.org/ |
| GraphPad Prism 9 | GraphPad Software | https://graphpad.com/scientific-software/prism/ |
| IgBlast | NCBI | https://www.ncbi.nlm.nih.gov/igblast/ |
| SnapGene | SnapGene | https://www.snapgene.com |
| MS Excel 2016 | Microsoft | https://www.microsoft.com/fr |
| Adobe Photoshop CC 2019 | Adobe | https://www.adobe.com/fr/products/photoshop/landpb.html |
| FB5P-seq | NA | https://github.com/MilpiedLab/FB5P-seq |
| Seurat v4 | NA | https://satijalab.org/seurat/ |

## METHODS

### Mouse model

*Aicda-Cre-ERT2 x Rosa26-lox-STOP-lox-eYFP* mice (20) were bred at the Centre d’Immuno-Phenomique, (Marseille, France), and transferred to the animal care facility of Centre d’Immunologie de Marseille-Luminy for experiments. All mice were maintained in the CIML mouse facility under specific pathogen-free conditions. Experimental procedures were conducted in agreement with French and European guidelines for animal care under the authorization number APAFIS #30945-2021040807508680, following review and approval by the local animal ethics committee in Marseille. Mice were used regardless of sex, at ages greater than 7 weeks and less than 3 months.

Mice were immunized with 100μg chicken ovalbumin (OVA) at 1μg/μl emulsified with Alum at a 1:1 (v:v) ratio, subcutaneously at the base of the tail, 50μl on each side. For induction of the Cre-ERT2-mediated labelling, we gavaged the mice once with 5mg of tamoxifen (TS648-1G, Sigma) in 200μL of peanut oil (P2144-250 ML, Sigma), at least 6 days after immunization. Mice were euthanized between 10 days and 21 days post-immunization (prime or boost) according to the experiment (**Figure 2B**).

**Figure 2.**
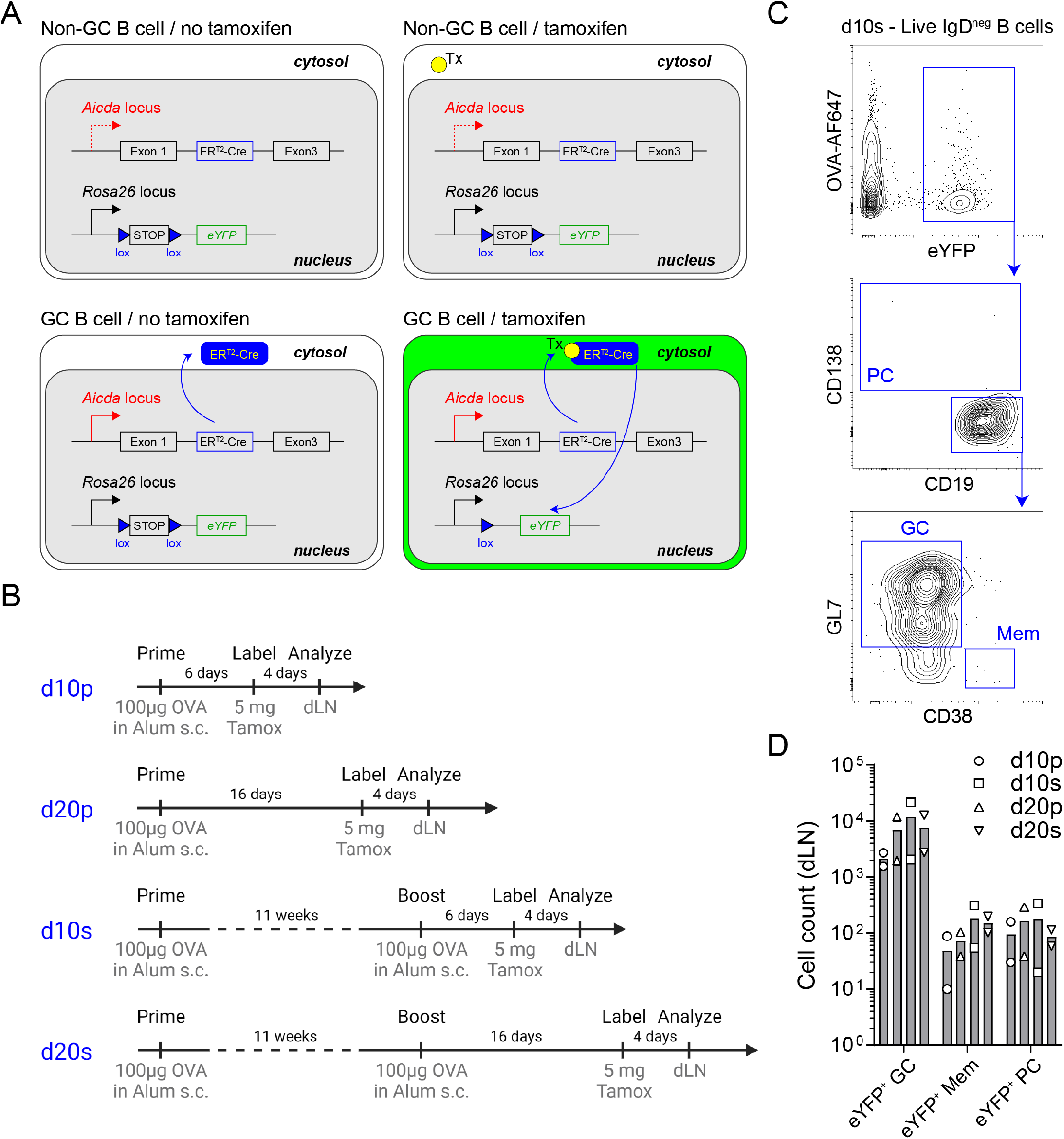
Experimental design for OVA-specific GC B cell analysis by FB5P-seq-mAbs. **A.** Schematic description of the Aicda-Cre-ERT2^+/-^ x Rosa26-eYFP-lox-stop-lox^+/-^ mouse model for fate mapping GC B cells and their progeny after tamoxifen gavage of immunized mice. **B**. Immunization strategies used to study GC B cells and their progeny at day 10 or day 20 after primary or secondary immunizations. **C**. Gating strategy for fate mapped GC B cells and their memory B cell (Mem) and plasma cell (PC) progeny. **D**. Absolute counts of eYFP^+^ GC B cells, eYFP^+^ Mem and eYFP^+^ PC in draining lymph nodes in d10p, d10s, d20p and d20s animals.

### Flow Cytometry and Cell Sorting of B Cell Subsets

Single-cell suspensions from draining lymph nodes were washed and resuspended in FACS buffer (5% fetal calf serum, 2mM EDTA, 5% Brilliant Stain Buffer Plus in PBS 1X) at a concentration of 100 million of cells per ml. Cells were first incubated with FcBlock (Biolegend) for 10 min on ice. Then, cells were incubated with a mix of antibodies (see table1 below) conjugated with fluorochromes 30 min on ice. Cells were washed in PBS, and incubated with the Live/Dead Fixable Aqua Dead Cell Stain (Thermofisher) for 10 min on ice. Cells were then washed again in FACS buffer and resuspended in 2% paraformaldehyde during 50min to preserve the eYFP contained in the cytoplasm. Cells were washed and permeabilized using the FoxP3 permeabilization kit (eBioscience) during 30min, then washed again in the permeabilization buffer and incubated with intracellular antibodies for 45min at RT. Cells were finally washed in permeabilization buffer and resuspended in FACS buffer. Cells were sorted on the BD Influx™ Cell Sorter, in 96-well plates, with index-sorting mode for recording the fluorescence parameters associated to each sorted cell.

### FB5P-seq library preparation, sequencing and data pre-processing

The protocol was performed as previously described by Attaf et al.(9). Individual cells were sorted into a 96-well PCR plate, with each well containing 2 μL of lysis buffer. Index sort mode was activated to record the fluorescence intensities of all markers for each individual cell. Flow cytometry standard (FCS) files from the index sort were analyzed using FlowJo software, and compensation parameters were exported as CSV tables for subsequent bioinformatic analysis. Immediately after sorting, plates containing individual cells were stored at -80°C until further processing. Following thawing, reverse transcription was performed, and the resulting cDNA was preamplified for 22 cycles. Libraries were then prepared according to the FB5P-seq protocol. The FB5P-seq data were processed to generate both a single-cell gene count matrix and single-cell B cell receptor (BCR) repertoire sequences for B cell analysis. Two separate bioinformatic pipelines were employed for gene expression and repertoire analysis, as detailed in Attaf et al.(9).

### Bioinformatics analysis

We used a custom bioinformatics pipeline to process fastq files and generate single-cell gene expression matrices and BCR sequence files as previously described(9). Detailed instructions for running the FB5P-seq bioinformatics pipeline can be found at https://github.com/MilpiedLab/FB5P-seq. Quality control was performed on each dataset independently to remove poor quality cells based on UMI counts, number of genes detected, ERCC spike-in quantification accuracy, and percentage of transcripts from mitochondrial genes. For each cell, gene expression UMI count values were log-normalized with Seurat *NormalizeData* with a scale factor of 10,000 to generate normalized UMI count matrices.

Index-sorting FCS files were visualized in FlowJo software and compensated parameters values were exported in CSV tables for further processing. For visualization on linear scales in the R programming software, we applied the hyperbolic arcsine transformation on fluorescence parameters(21).

For BCR sequence reconstruction, the outputs of the FB5P-seq pipeline were further processed and filtered with custom R scripts. For each cell, reconstructed contigs corresponding to the same V(D)J rearrangement were merged, keeping the largest sequence for further analysis. We discarded contigs with no constant region identified in-frame with the V(D)J rearrangement. In cases where several contigs corresponding to the same BCR chain had passed the above filters, we retained the contig with the highest expression level. BCR metadata from the *MigMap* and *Blastn* annotations were appended to the gene expression and index sorting metadata for each cell.

Supervised annotation of scRNA-seq datasets were performed as described extensively in **Figure 2B** and Supplementary **Figure 2C** of Binet et al.(22). Briefly, we used the *AddModuleScore* function to compute gene expression scores for every cell in the dataset for the cell type specific signatures. For DZ and LZ signatures, genes associated to cell cycle ontologies (based on GO terms), were removed from the gene lists prior to scoring, as described in Milpied et al.(23). Thresholds for “gating” were defined empirically. Single-cell gene expression heatmaps were generated by the *doheatmap* function in *R*.

### BCR amplification and cloning

From the FACS-sorted single-cell RNA-seq libraries in 96-well PCR plates, 2 μl of each well of the plate was diluted. This diluted cDNA was used to amplify the variable regions with a multiplex PCR (**Supplementary Table 1**). After verification of the amplification on an agarose gel, the PCR products were purified and adjusted to the concentration necessary to perform the cloning. Cloning of the variable regions was done by SLIC (Sequence and ligation-independent cloning) in each linearized and pre-purified expression vector (24). To verify that the inserts had been cloned into each expression vector, the colonies were screened by PCR and a bacterial colony fingerprint was made. The primers used were specific to each IgH and IgK vector (**Supplementary Table 2**).

Plasmid DNA preparations were made from the screened positive bacterial fingerprints and each construct was sequenced and checked against the original sequences.

### Antibody production

The Expi293™ cells grown in Expi293™ Expression Medium were cultured in sterile vented-cap Erlenmeyer flasks. On the day of transfection, the heavy chain-containing vectors and light chain-containing vectors were incubated in the presence of polyethyleneimine. The cells were incubated for 6 days in the incubator. The culture supernatants were then centrifuged, filtered and the antibodies present in supernatant were purified on protein G beads on polyprep chromatography columns.

### Antibody analysis by ELISA

OVA antigen was coated in 96-well ELISA plates for incubation overnight at 4°C. After 3 washes, the plate was blocked with 100 μl of blocking buffer and incubated for 2h at room temperature. After 3 washes, decreasing concentrations of control and target antibodies were added and incubated for 2h at room temperature. After 3 washes, HRP-coupled anti-mouse IgG secondary antibody diluted 4000-fold was added and incubated for 2h at room temperature. After 3 washes, 100 μl of TMB solution was added and incubated for 15 min. The reaction was stopped by adding 100 μl of 2N sulfuric acid. The reaction was measured with a plate reader at 450 nm.

### Detailed protocol

A detailed reagent setup protocol is presented in Supplementary File 1.

A detailed step-by-step protocol is presented in Supplementary File 2.

## LIMITATIONS

The amplification of the Ig genes may be dependent on the sequence of the V genes sequences of the isolated B cells. It may be necessary to amplify the BCR from a larger number of cells to obtain the sufficient number of Ig. However, the knowledge of each BCR sequence upstream of the amplification, can be a precious help in the failure of some V gene PCR products, by being able to specifically adapt the sequence of the primers.

The cloning of variable chains by the SLIC method can be variable. This can be due to the purification of the PCR products or the efficiency of the reaction. In this case, it is possible to overcome this problem by a classical cloning technique which is made possible by the presence of restriction enzyme sites in the PCR products of each chain.

The Elisa control experiment which is performed the day before antibody purification is a good indication of antibody production in the culture supernatant. However, the level of production does not always reflect the total amount of antibodies obtained after purification. This depends on the ability of the antibodies to bind to the protein-G beads due to their structure or purity.

## TROUBLESHOOTING

A detailed troubleshooting table is presented as Supplementary File 3.

## RESULTS

We applied FB5P-seq-mAbs to study murine GC B cells after prime or prime-boost immunization with the model antigen chicken ovalbumin (OVA). We used the Aicda-Cre-ERT2^+/-^ x Rosa26-eYFP-lox-stop-lox^+/-^ mouse model (20,25) for fate mapping GC B cells and their progeny after tamoxifen gavage of immunized mice (**Figure 2A**). Mice were immunized with OVA and alum adjuvant subcutaneously, once for prime immunizations, a second time 11 weeks later for prime-boost immunizations, and gavaged with tamoxifen 4 days before sacrifice and collection of draining lymph nodes; draining lymph nodes were collected 10 or 20 days after primary (d10p, d20p), or after secondary (d10s, d20s) immunization (**Figure 2B**). GC B cells and their progeny were gated as live IgD-negative B cells expressing eYFP, and were further subdivided as CD138^+^ plasma cells (PC), CD19^+^GL7^+^CD38^-^ GC B cells, or CD19^+^CD38^+^GL7^-^ memory B cells (Mem) (**Figure 2C**). As expected, eYFP^+^ GC B cells outnumbered their Mem and PC progeny at days 10 or 20 after primary or secondary immunization in draining lymph nodes (**Figure 2D**).

We sorted single IgD^neg^ eYFP^+^ B cells from d10p, d20p, d10s and d20s draining lymph nodes and prepared 5’-end scRNA-seq libraries with the FB5P-seq protocol. All surface staining parameters, including binding of OVA-AlexaFluor647 (OVA-AF647) antigen, were recorded by index sorting and appended to the gene-by-cell UMI count matrix. We also used the FB5P-seq bioinformatic pipeline to reconstruct the *Igh* and *Igκ/λ* variable region sequences from 5’-end scRNA-seq reads (9). After quality controls, we retained 769 cells, including 573 cells with paired full *Igh* and *Igκ/λ* variable region sequences. Based on signature gene expression, we annotated the majority of cells as LZ (n=331), DZ (n=300) or recycling LZtoDZ (n=55) GC B cells, and identified minor fractions of putative preMem (n=29), prePC (n=27), or *bona fide* PC (n=5) (**Figure 3A**). Consistent with other scRNA-seq studies of murine GC B cells (26–28), LZ cells expressed high levels of *Fcer2a* (encoding CD23), *Cd83, H2-Oa* (encoding MHC-II subunit), *Cd86* and *Nfkbia* transcripts and were mostly quiescent or in S phase. DZ cells expressed high levels of *Gcsam, Ccnb2, Hmces, Pafah1b3* and *Stmn1* transcripts, and were mostly in G2/M phase. LZtoDZ cells expressed both LZ and DZ marker genes, as well as high levels of *C1qbp, Pa2g4, Apex1, Nop58* and *Exosc7* transcripts, and were mostly in S phase. PreMem cells expressed LZ marker genes, as well as high levels of *Serpinb1a, Capg, Ms4a4c, Gpr183* and *Btg1* transcripts, and were quiescent. PrePC cells were transcriptionally close to LZtoDZ cells, with additional expression of *Sub1, Pdia4, Emb, Glo1, Polr2h* transcripts, and were mostly proliferating. PC expressed high levels of some PrePC marker genes (*Sub1, Pdia4*) and other transcripts associated with antibody-producing cells differentiation such as *Sdc1* (encoding CD138), *Fam46c, Irf4, Bst2*, and *Glipr1*.

**Figure 3.**
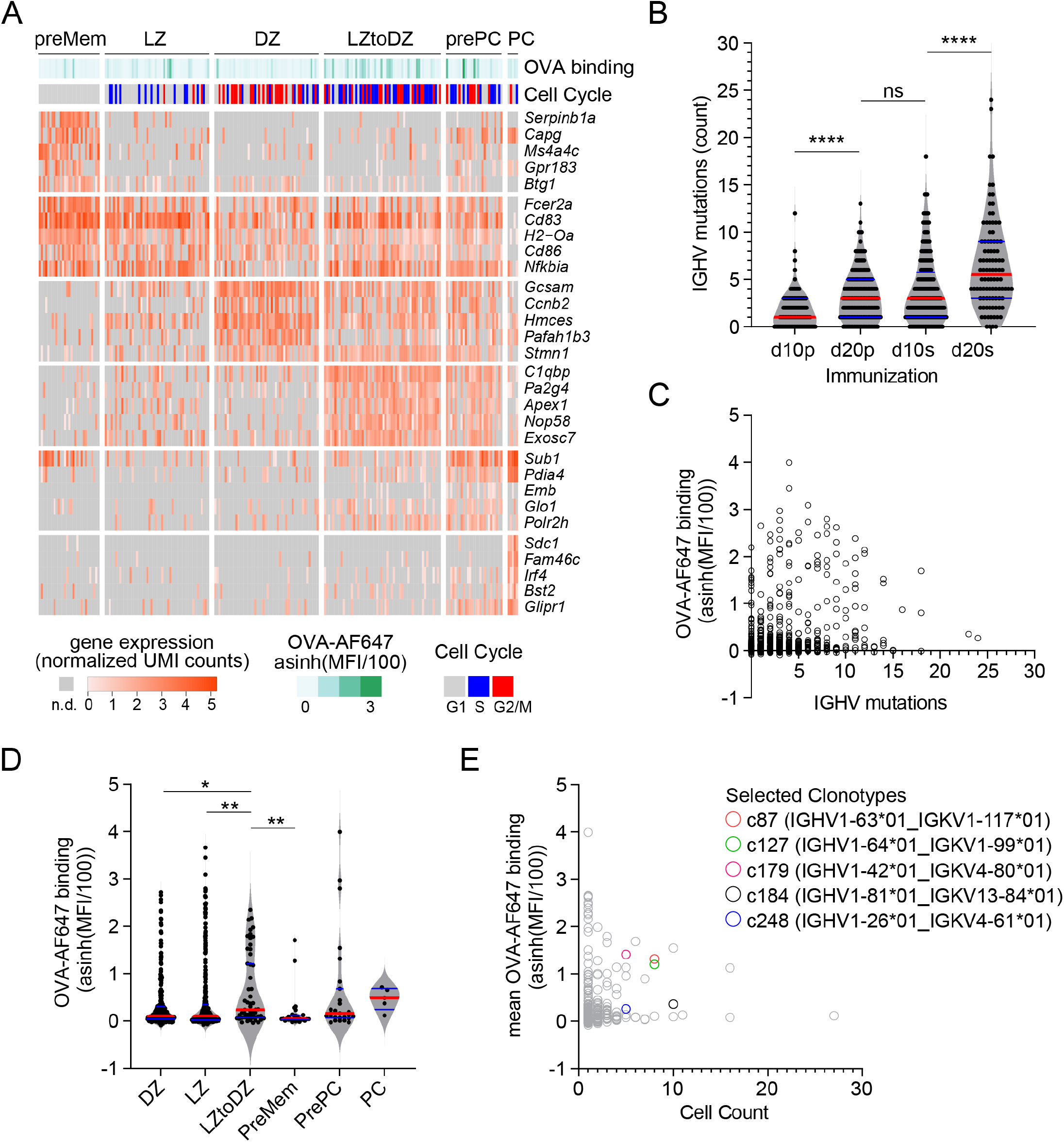
FB5P-seq analysis of OVA-specific GC B cells. **A.** Gene expression heatmap showing expression of top 5 marker genes for annotated subsets of GC B cells. For visualization, only randomly sampled 50 cells from LZ and DZ subsets are displayed on the heatmap. Bars above the heatmap indicate OVA-AF647 surface binding (fluorescence intensity) and cell cycle phase inferred from gene expression. **B**. Number of mutations in *Ighv* sequences of cells from d10p, d20p, d10s, and d20s samples, as compared to their inferred germline. Each dot is a cell. ****: p<0.0001, ns: not significant, adjusted p-value for Dunn’s multiple comparison test after significant Kruskal-Wallis analysis. **C**. Scatter plot of *Ighv* mutation count (x-axis) by OVA-AF647 surface binding intensity (y-axis). Each dot is a cell. **D**. OVA-AF647 surface binding intensity in cells from different gene expression-based subsets. Each dot is a cell. *: p<0.05, **: p<0.01, adjusted p-value for Dunn’s multiple comparison test after significant Kruskal-Wallis analysis. **E**. Scatter plot of clonotype size (x-axis) by mean OVA-AF647 surface binding intensity (y-axis) for all clonotypes. Each dot is a clonotype, colored symbols for clonotypes selected for further analyses.

Consistent with ongoing affinity maturation occurring in primary and secondary GC, the number of *Ighv* mutations increased from d10p to d20p, and then further increased from d10s to d20s (**Figure 3B**). Surface binding of OVA-AF647 antigen, a flow cytometry-based measurement of BCR affinity for the immunizing OVA antigen, was not correlated to the number of *Ighv* mutations (**Figure 3C**). Among the different cell subsets, recycling LZtoDZ cells had the highest rate of OVA-AF647 binding (**Figure 3D**), consistent with high affinity antigen-specific cells being selected and recycled from LZ GC B cells at all stages of primary and secondary GC reactions. We used *Ighv* sequence information to identify groups of clonally related cells (clonotypes), and selected 5 clonotypes of size ≥5 containing at least 1 cell with significant OVA-AF647 surface binding (**Figure 3E**) for recombinant mAb expression.

For each of those selected clonotypes, we designed unmutated germline sequences for the *Ighv* and *Igkv* chains, and had them synthesized with appropriate 5’ and 3’ ends for direct cloning into our IgG1 and IgK expression vectors. All other *Ighv* and *Igkv* sequences were amplified and cloned directly from the archived single-cell cDNA obtained in the FB5P-seq protocol, as detailed in the methods. We succeeded to obtain significant amounts of recombinant mAb for 5/5 cells for clonotype c179, 5/5 cells for clonotype c248, 7/8 cells for clonotype c127, 7/8 cells for clonotype c87, and 8/10 cells for clonotype c184. Failures were due to unsuccessful PCR for one of the two chains of a given cell. In some cases where the multiplexed PCR failed, we used the reconstructed *Ighv* sequence from the FB5P-seq pipeline to design sequence-specific forward primers for *Ighv* amplification, and obtained PCR products that were cloned and used to produce mAbs.

We then tested serial dilutions of those mAbs for OVA binding in indirect ELISA assays (**Figure 4A-E**). For all clonotypes, at least one mAb detectably bound OVA, albeit at very high concentration for clonotype c248 (**Figure 4B**). As expected, there was intraclonal heterogeneity in antigen binding capacity of single-cell mAbs, with most mAbs binding better than the germline for all clonotypes except c127 (**Figure 4A-E**), but some mAbs having no detectable OVA binding for clonotypes c179 (**Figure 4A**), c248 (**Figure 4B**), c87 (**Figure 4D**) and c184 (**Figure 4E**). For clonotype c127, none of the GC B cell-derived mAbs bound better than the germline (**Figure 4C**). Based on the ELISA binding curves, we computed the ELISA Binding Threshold values for all mAbs, reflecting the minimum amount of mAbs giving detectable binding in our ELISA assays. There was no association between the number of mutations in the *Ighv* or *Igkv* regions and the ELISA Binding Threshold (**Table 1**), but there was a significant correlation between the ELISA Binding Threshold and the OVA-AF647 surface binding measured by index sorting (**Table 1** and **Figure 5A**), even for intraclonal comparisons. Most cells which had surface binding of OVA-AF647 above 1, corresponding to signal above background noise, produced mAbs with ELISA Binding Threshold below 10 ng. Conversely, most cells which had surface binding of OVA-AF647 below 1 produced mAbs with ELISA Binding Threshold above 100 ng. Thus, OVA-AF647 surface binding is a good proxy for assessing antigen-binding properties of the BCR in single GC B cells. Nevertheless, 5 outlier cells, 3 of which were from clonotype c184, had no detectable surface binding of OVA-AF647, but produced mAbs with ELISA Binding Threshold values between 10 and 100 ng, suggesting that the FACS assay may be less sensitive than the ELISA on recombinant mAbs. We inspected the expression of subset-specific marker genes in individual cells in relation with the binding capacities of their BCR measured by FACS and ELISA (**Figure 5B**). Consistent with our observations on the total dataset (**Figure 3D**), it was interesting to note that for clonotypes c179, c87 and c127, most of the cells with good OVA-binding capacity expressed genes associated with LZ, DZ, and LZtoDZ states. For both clonotypes c127 and c184, we captured one cell in PreMem state that expressed OVA-binding BCR with ELISA Binding Threshold values in the 10 ng range, consistent with the selection of Mem B cells from mid-affinity GC B cells (29).

**Table 1.** Characteristics of recombinant mAbs produced from OVA-specific GC B cell clonotypes. For all mAbs grouped by clonotype, the number of mutations in the CDR or framework (FW) regions of the heavy (VH) and light (VK) chains are indicated in the first 4 columns, the ELISA Binding Threshold in ng in the 5^th^ column, and the OVA-AF647 surface binding measured by index sorting in the 6^th^ column.

|  | number of mutations |  |  |  | ELISA binding<br>threshold (ng) | OVA-AF647 binding<br>(asinh(MFI/100)) |
| --- | --- | --- | --- | --- | --- | --- |
|  | VH |  | VK |  |  |  |
|  | CDR | FW | CDR | FW |  |  |
| **c179 (IGHV1-42\*01_IGKV4-80\*01)** |  |  |  |  |  |  |
| germline | 0 | 0 | 0 | 0 | >1000 | NA |
| p7.BC76 | 1 | 0 | 1 | 0 | 19.03 | 2.14 |
| p7.BC41 | 0 | 1 | 2 | 1 | >1000 | 0.06 |
| p7.BC49 | 0 | 1 | 0 | 0 | 0.27 | 1.86 |
| p8.BC75 | 1 | 0 | 1 | 0 | 4.85 | 1.53 |
| p8.BC46 | 0 | 0 | 0 | 0 | 3.86 | 1.46 |
| **c248 (IGHV1-26\*01_IGKV4-61\*01)** |  |  |  |  |  |  |
| germline | 0 | 0 | 0 | 0 | >1000 | NA |
| p9.BC45 | 2 | 3 | 0 | 0 | >1000 | 0.03 |
| p9.BC76 | 2 | 1 | 0 | 0 | 14.94 | 0.04 |
| p9.BC37 | 2 | 1 | 0 | 0 | 615.18 | 0.44 |
| p9.BC61 | 0 | 1 | 0 | 0 | 539 | 0.21 |
| p9.BC64 | 0 | 0 | 0 | 0 | 412 | 0.60 |
| **c127 (IGHV1-64\*01_IGKV1-99\*01)** |  |  |  |  |  |  |
| germline | 0 | 0 | 0 | 0 | 0.75 | NA |
| p9.BC16 | 1 | 1 | 0 | 2 | 219 | 1.35 |
| p9.BC78 | 1 | 0 | 0 | 0 | 502 | 0.27 |
| p9.BC41 | 1 | 1 | 0 | 1 | 1.74 | 1.89 |
| p10.BC3 | 0 | 0 | 0 | 0 | 1.6 | 1.57 |
| p9.BC1 | 1 | 0 | 0 | 0 | 129 | 0.68 |
| p9.BC18 | 0 | 0 | 0 | 0 | 3.2 | 1.27 |
| p10.BC50 | 1 | 1 | 0 | 0 | 8.9 | 2.06 |
| **c87 (IGHV1-63\*01_IGKV1-117\*01)** |  |  |  |  |  |  |
| germline | 0 | 0 | 0 | 0 | >1000 | NA |
| p8.BC50 | 2 | 5 | 2 | 3 | 0.8 | 1.93 |
| p7.BC31 | 2 | 1 | 0 | 1 | 336 | 0.48 |
| p7.BC19 | 1 | 2 | 0 | 2 | 10.8 | 0.24 |
| p7.BC16 | 1 | 3 | 1 | 0 | 1.94 | 1.43 |
| p8.BC83 | 2 | 1 | 1 | 1 | 1.73 | 1.41 |
| p7.BC1 | 0 | 0 | 1 | 0 | 0.74 | 1.64 |
| p7.BC27 | 0 | 0 | 0 | 0 | 0.42 | 1.09 |
| **c184 (IGHV1-81\*01_IGKV13-84\*01)** |  |  |  |  |  |  |
| germline | 0 | 0 | 0 | 0 | >1000 | NA |
| p10.BC30 | 1 | 1 | 0 | 4 | 14.6 | 0.13 |
| p9.BC6 | 2 | 0 | 2 | 2 | >1000 | 0.03 |
| p10.BC57 | 0 | 0 | 0 | 0 | 352 | 1.47 |
| p9.BC75 | 1 | 1 | 0 | 0 | 451 | 0.05 |
| p10.BC74 | 1 | 0 | 0 | 4 | 24.8 | 0.33 |
| p10.BC38 | 1 | 1 | 0 | 0 | 280 | 0.94 |
| p9.BC25 | 1 | 2 | 0 | 0 | >1000 | 0.00 |
| p9.BC40 | 2 | 0 | 0 | 0 | 17 | 0.31 |

**Figure 4.**
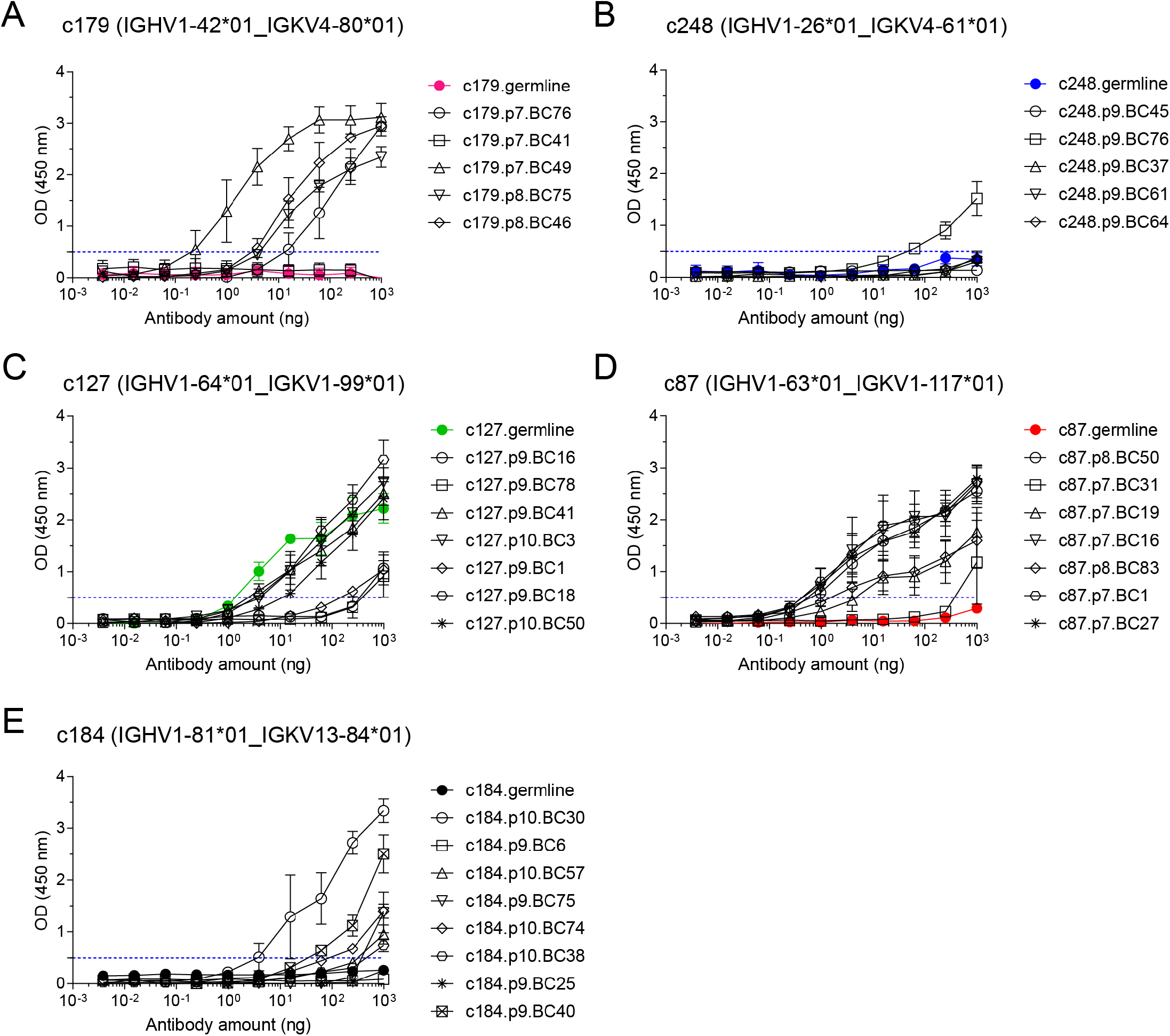
OVA binding capacity of recombinant mAbs measured by indirect ELISA assay. Recombinant mAbs from clonotypes c179 (**A**), c248 (**B**), c127 (**C**), c87 (**D**), and c184 (**E**) and their respective germline (colored line and symbols in each plot) were tested as serial 4-fold dilutions (starting from 1μg per well) in indirect ELISA on OVA-coated plates. All measurements were performed independently from 2 to 5 times, and mean ± s.e.m. OD values are reported as connected symbols. The dotted blue line represents the threshold OD value that was used to compute the ELISA Binding Threshold value for each mAb.

**Figure 5.**
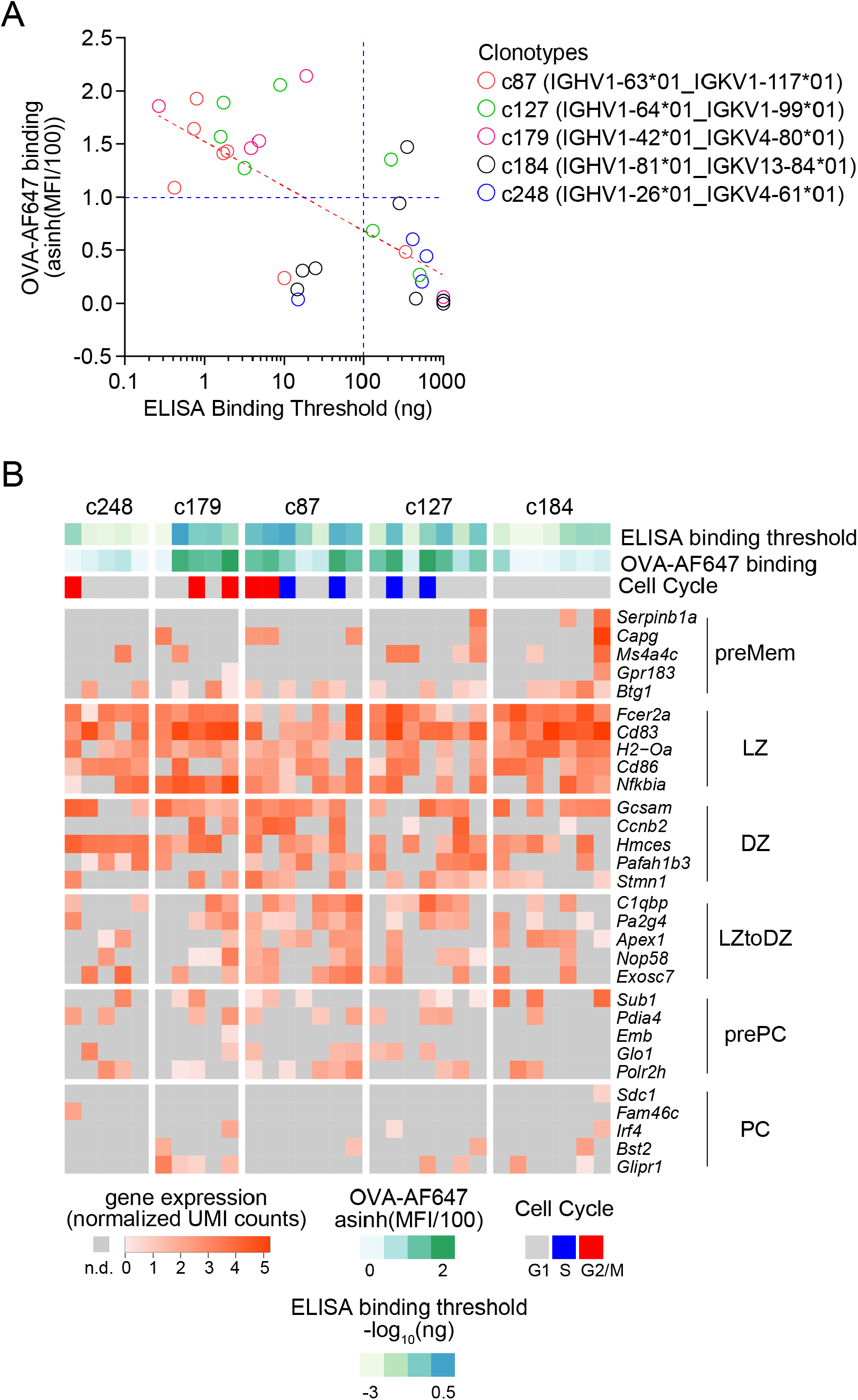
Integrative analysis of antigen-binding and gene expression of OVA-specific GC B cells. **A**. Scatter plot of ELISA Binding Threshold (x-axis) by OVA-AF647 binding (y-axis) for mAbs of the indicated clonotypes (color key). The dashed red line indicates the non-linear semi-log least-squares fit (R^2^= 0.4638). The dashed blue lines are manually set thresholds to define good OVA binders measured by ELISA (x-axis) or FACS (y-axis). **B**. Gene expression heatmap showing expression of top 5 marker genes of subsets of GC B cells (as in **Figure 3A**) in cells from the selected clonotypes studied by ELISA. Bars above the heatmap indicate OVA ELISA Binding Threshold, OVA-AF647 surface binding (fluorescence intensity) and cell cycle phase inferred from gene expression.

Altogether, those results illustrate some of the quantitative single-cell analyses that are made possible when integrating phenotypic, transcriptomic, molecular and biochemical readouts on single B cells with FB5P-seq-mAbs.

## DISCUSSION

FB5P-seq-mAbs was designed as an extension of our plate-based 5’-end scRNA-seq method, FB5P-seq (9). As such, it can be performed on archived single-cell cDNA months to years after preparation of the initial FB5P-seq libraries. Here, we demonstrated FB5P-seq-mAbs on mouse OVA-specific GC B cells, but it could also be applied to other antigen-specific B cell types. For example, we recently applied FB5P-seq to characterize memory B cells in mouse lungs after influenza virus infection and discovered bystander Mem B cells with no apparent specificity to influenza virus antigens (11). In that case, FB5P-seq-mAbs may be useful to produce recombinant mAbs from bystander Mem B cells and screen those mAbs for other (auto)antigen specificities.

The FB5P-seq-mAbs protocol can be easily adapted to study human B cells by using the *IGH* and *IGK/L* amplification and cloning strategies described by others previously (13). In FB5P-seq-mAbs, the scRNA-seq and BCR sequence reconstruction are performed before cloning of *IGH* and *IGK/L*, giving the opportunity to select only the single-cell cDNAs from specific cell states and/or clonotypes as starting material for mAb production. Knowing the *IGH* and *IGK/L* sequences before cloning is also particularly interesting when working with highly mutated B cells, because it allows the design of cell-specific or clonotype-specific PCR primers instead of multiplexed PCR primers that may not be optimal.

FB5P-seq-mAbs is modular, and equivalent integrative single-cell analyses of B cells may be obtained via distinct alternatives. Other plate-based single-cell RNA-seq library preparation protocols, such as Smart-seq2 (7) or Smart-seq3 (30), may be used to produce single-cell gene expression and BCR sequencing data and archive single-cell cDNA for recombinant mAb production. Other cloning and expression methods may be used to produce recombinant mAbs from single-cell cDNA(31). Another possibility is to use high-throughput droplet-based methods for characterization of phenotype, gene expression, BCR sequence and antigen binding (12), then select cells of interest *in silico* and have their *IGH* and *IGK/L* V genes synthesized for direct cloning and production.

The two main directions for improving FB5P-seq-mAbs are depth and throughput. First, we may modify the FB5P-seq protocol to adopt recent improvements to plate-based scRNA-seq protocols that make them more sensitive and easier to perform. For example, we may implement the one-step RT-PCR of the recently published FLASH-seq-UMI protocol (32) which resulted in higher sensitivity and shorter library preparation time when compared to the Smart-seq3 approach. Second, we may implement antibody cloning and production strategies that are designed for higher throughput (33,34) and can be automatized.

FB5P-seq-mAbs and its future improved versions will be important to continue the in-depth studies of antigen-specific B cell responses in animal models, and to discover the antigen reactivity and affinity of B cells in human infectious diseases, autoimmunity and cancer.

## ACKNOWLEDGEMENTS

We thank all members of the Integrative B cell Immunology lab of CIML for useful comments and discussions. We thank the Animal Care, Flow Cytometry, Genomics, and Computational Biology, Bioinformatics and Modeling core facilities of CIML for support in our experiments and analyses. We thank Dr Lotta von Boehmer and Anna Gazumyan from Pr Michel Nussenzweig’s lab for sending us antibody production vectors and providing useful tips for antibody production. We thank Dr Benjamin Rossi from Innate Pharma for providing useful tips for antibody production. Centre de Calcul Intensif d’Aix-Marseille is acknowledged for granting access to its high performance computing resources. This work was supported by grants from ANR (ANR-17-CE15-0009-01 JCJC MoDEx-GC) and Inserm ITMO Cancer (grant number ASC19008ASA) to P.M. This work was supported by institutional grants from INSERM, CNRS, and Aix-Marseille University to the CIML. N.A. was supported by fellowships from the French Ministry of Research and Higher Education.

## AUTHOR CONTRIBUTIONS

S.A. prepared the detailed protocol and wrote the manuscript; N.A. performed the OVA immunization experiments and FB5P-seq library preparations; C.D. performed bioinformatics analyses; M.M. set up the OVA-binding ELISA assay; A.C. contributed to the design of the antibody cloning and production protocol; P.M. supervised all experiments, acquired funding, analyzed the data, prepared figures and wrote the manuscript; J-M.N. developed the protocol, performed mAb cloning, production and *in vitro* assays, analyzed the data, prepared figures and wrote the manuscript.

## REFERENCES

1. McHeyzer-Williams M, Okitsu S, Wang N, McHeyzer-Williams L. Molecular programming of B cell memory. Nat Rev Immunol (2012) 12:24–34. doi: 10.1038/nri3128

2. Cyster JG, Allen CDC. B Cell Responses: Cell Interaction Dynamics and Decisions. Cell (2019) 177:524–540. doi: 10.1016/j.cell.2019.03.016

3. Stavnezer J, Guikema JEJ, Schrader CE. Mechanism and Regulation of Class Switch Recombination. Annual Review of Immunology (2008) 26:261–292. doi: 10.1146/annurev.immunol.26.021607.090248

4. Peled JU, Kuang FL, Iglesias-Ussel MD, Roa S, Kalis SL, Goodman MF, Scharff MD. The Biochemistry of Somatic Hypermutation. Annual Review of Immunology (2008) 26:481–511. doi: 10.1146/annurev.immunol.26.021607.090236

5. Victora GD, Nussenzweig MC. Germinal centers. Annu Rev Immunol (2012) 30:429–457. doi: 10.1146/annurev-immunol-020711-075032

6. Attaf N, Baaklini S, Binet L, Milpied P. Heterogeneity of germinal center B cells: New insights from single-cell studies. European Journal of Immunology n/a: doi: 10.1002/eji.202149235

7. Picelli S, Björklund ÅK, Faridani OR, Sagasser S, Winberg G, Sandberg R. Smart-seq2 for sensitive full-length transcriptome profiling in single cells. Nature Methods (2013) 10:1096–1098. doi: 10.1038/nmeth.2639

8. Lindeman I, Emerton G, Mamanova L, Snir O, Polanski K, Qiao S-W, Sollid LM, Teichmann SA, Stubbington MJT. BraCeR: B-cell-receptor reconstruction and clonality inference from single-cell RNA-seq. Nat Methods (2018) 15:563–565. doi: 10.1038/s41592-018-0082-3

9. Attaf N, Cervera-Marzal I, Dong C, Gil L, Renand A, Spinelli L, Milpied P. FB5P-seq: FACS-Based 5-Prime End Single-Cell RNA-seq for Integrative Analysis of Transcriptome and Antigen Receptor Repertoire in B and T Cells. Front Immunol (2020) 11: doi: 10.3389/fimmu.2020.00216

10. Mimitou EP, Cheng A, Montalbano A, Hao S, Stoeckius M, Legut M, Roush T, Herrera A, Papalexi E, Ouyang Z, et al. Expanding the CITE-seq tool-kit: Detection of proteins, transcriptomes, clonotypes and CRISPR perturbations with multiplexing, in a single assay. Nat Methods (2019) 16:409–412. doi: 10.1038/s41592-019-0392-0

11. Gregoire C, Spinelli L, Villazala-Merino S, Gil L, Holgado MP, Moussa M, Dong C, Zarubica A, Fallet M, Navarro J-M, et al. Viral infection engenders bona fide and bystander subsets of lung-resident memory B cells through a permissive mechanism. Immunity (2022) doi: 10.1016/j.immuni.2022.06.002

12. Setliff I, Shiakolas AR, Pilewski KA, Murji AA, Mapengo RE, Janowska K, Richardson S, Oosthuysen C, Raju N, Ronsard L, et al. High-Throughput Mapping of B Cell Receptor Sequences to Antigen Specificity. Cell (2019) 179:1636-1646.e15. doi: 10.1016/j.cell.2019.11.003

13. Tiller T, Meffre E, Yurasov S, Tsuiji M, Nussenzweig MC, Wardemann H. Efficient generation of monoclonal antibodies from single human B cells by single cell RT-PCR and expression vector cloning. J Immunol Methods (2008) 329:112–124. doi: 10.1016/j.jim.2007.09.017

14. Tiller T, Busse CE, Wardemann H. Cloning and expression of murine Ig genes from single B cells. Journal of Immunological Methods (2009) 350:183–193. doi: 10.1016/j.jim.2009.08.009

15. von Boehmer L, Liu C, Ackerman S, Gitlin AD, Wang Q, Gazumyan A, Nussenzweig MC. Sequencing and cloning of antigen-specific antibodies from mouse memory B cells. Nat Protocols (2016) 11:1908– 1923. doi: 10.1038/nprot.2016.102

16. Klein F, Mouquet H, Dosenovic P, Scheid J, Scharf L, Nussenzweig MC. Antibodies in HIV-1 Vaccine Development and Therapy. Science (2013) 341:1199–1204. doi: 10.1126/science.1241144

17. Vanshylla K, Fan C, Wunsch M, Poopalasingam N, Meijers M, Kreer C, Kleipass F, Ruchnewitz D, Ercanoglu MS, Gruell H, et al. Discovery of ultrapotent broadly neutralizing antibodies from SARS-CoV-2 elite neutralizers. Cell Host & Microbe (2022) 30:69-82.e10. doi: 10.1016/j.chom.2021.12.010

18. Viant C, Weymar GHJ, Escolano A, Chen S, Hartweger H, Cipolla M, Gazumyan A, Nussenzweig MC. Antibody Affinity Shapes the Choice between Memory and Germinal Center B Cell Fates. Cell (2020) doi: 10.1016/j.cell.2020.09.063

19. Mayer CT, Gazumyan A, Kara EE, Gitlin AD, Golijanin J, Viant C, Pai J, Oliveira TY, Wang Q, Escolano A, et al. The microanatomic segregation of selection by apoptosis in the germinal center. Science (2017)eaao2602. doi: 10.1126/science.aao2602

20. Le Gallou S, Nojima T, Kitamura D, Weill J-C, Reynaud C-A. The AID-Cre-ERT2 Model: A Tool for Monitoring B Cell Immune Responses and Generating Selective Hybridomas. Methods Mol Biol (2017) 1623:243–251. doi: 10.1007/978-1-4939-7095-7_19

21. Finak G, Perez J-M, Weng A, Gottardo R. Optimizing transformations for automated, high throughput analysis of flow cytometry data. BMC Bioinformatics (2010) 11:546. doi: 10.1186/1471-2105-11-546

22. Binet L, Dong C, Attaf N, Gil L, Fallet M, Boudier T, Escalière B, Chasson L, Siret C, Pavert SA van de, et al. Specific pre-plasma cell states and local proliferation at the dark zone – medulla interface characterize germinal center-derived plasma cell differentiation in lymph node. (2024)2024.07.26.605240. doi: 10.1101/2024.07.26.605240

23. Milpied P, Cervera-Marzal I, Mollichella M-L, Tesson B, Brisou G, Traverse-Glehen A, Salles G, Spinelli L, Nadel B. Human germinal center transcriptional programs are de-synchronized in B cell lymphoma. Nat Immunol (2018) 19:1013–1024. doi: 10.1038/s41590-018-0181-4

24. Jeong J-Y, Yim H-S, Ryu J-Y, Lee HS, Lee J-H, Seen D-S, Kang SG. One-Step Sequence- and Ligation-Independent Cloning as a Rapid and Versatile Cloning Method for Functional Genomics Studies. Appl Environ Microbiol (2012) 78:5440–5443. doi: 10.1128/AEM.00844-12

25. Dogan I, Bertocci B, Vilmont V, Delbos F, Mégret J, Storck S, Reynaud C-A, Weill J-C. Multiple layers of B cell memory with different effector functions. Nat Immunol (2009) 10:1292–1299. doi: 10.1038/ni.1814

26. Kennedy DE, Okoreeh MK, Maienschein-Cline M, Ai J, Veselits M, McLean KC, Dhungana Y, Wang H, Peng J, Chi H, et al. Novel specialized cell state and spatial compartments within the germinal center. Nature Immunology (2020)1–11. doi: 10.1038/s41590-020-0660-2

27. Pikor NB, Mörbe U, Lütge M, Gil-Cruz C, Perez-Shibayama C, Novkovic M, Cheng H-W, Nombela-Arrieta C, Nagasawa T, Linterman MA, et al. Remodeling of light and dark zone follicular dendritic cells governs germinal center responses. Nature Immunology (2020)1–11. doi: 10.1038/s41590-020-0672-y

28. Nakagawa R, Toboso-Navasa A, Schips M, Young G, Bhaw-Rosun L, Llorian-Sopena M, Chakravarty P, Sesay AK, Kassiotis G, Meyer-Hermann M, et al. Permissive selection followed by affinity-based proliferation of GC light zone B cells dictates cell fate and ensures clonal breadth. PNAS (2021) 118: doi: 10.1073/pnas.2016425118

29. Shinnakasu R, Inoue T, Kometani K, Moriyama S, Adachi Y, Nakayama M, Takahashi Y, Fukuyama H, Okada T, Kurosaki T. Regulated selection of germinal-center cells into the memory B cell compartment. Nature Immunology (2016) 17:861–869. doi: 10.1038/ni.3460

30. Hagemann-Jensen M, Ziegenhain C, Chen P, Ramsköld D, Hendriks G-J, Larsson AJM, Faridani OR, Sandberg R. Single-cell RNA counting at allele and isoform resolution using Smart-seq3. Nature Biotechnology (2020) 38:708–714. doi: 10.1038/s41587-020-0497-0

31. Ho IY, Bunker JJ, Erickson SA, Neu KE, Huang M, Cortese M, Pulendran B, Wilson PC. Refined protocol for generating monoclonal antibodies from single human and murine B cells. J Immunol Methods (2016) 438:67–70. doi: 10.1016/j.jim.2016.09.001

32. Hahaut V, Pavlinic D, Carbone W, Schuierer S, Balmer P, Quinodoz M, Renner M, Roma G, Cowan CS, Picelli S. Fast and highly sensitive full-length single-cell RNA sequencing using FLASH-seq. Nat Biotechnol (2022)1–5. doi: 10.1038/s41587-022-01312-3

33. Gieselmann L, Kreer C, Ercanoglu MS, Lehnen N, Zehner M, Schommers P, Potthoff J, Gruell H, Klein F. Effective high-throughput isolation of fully human antibodies targeting infectious pathogens. Nat Protoc (2021) 16:3639–3671. doi: 10.1038/s41596-021-00554-w

34. Han X, Wang Y, Li S, Hu C, Li T, Gu C, Wang K, Shen M, Wang J, Hu J, et al. A Rapid and Efficient Screening System for Neutralizing Antibodies and Its Application for SARS-CoV-2. Front Immunol (2021) 12:653189. doi: 10.3389/fimmu.2021.653189

