## Supplementary Table 1 for "FB5P-seq-mAbs: monoclonal antibody production from FB5P-seq libraries for integrative single-cell analysis of B cells"

**Supplementary Table 1: Primers for PCR amplification of the heavy and light chain genes**

The restriction enzyme sites are underlined. The additional nucleotides in the reverse IgG1 primers are in bold.

**Multiplex forward (for mouse IgG1 expression vector)**

| T4hmFH_I | CTAGTAGCAACTGCAACCGGTGTACATTCTGAWGTGCAGCTGGTGGAGTC |
| --- | --- |
| T4hmFH_II | CTAGTAGCAACTGCAACCGGTGTACATTCTCAGGTGCAGCTGAAGSAGTC |
| T4hmFH_III | CTAGTAGCAACTGCAACCGGTGTACATTCTGARGTGAAGCTGGTGGARTC |
| T4hmFH_IV | CTAGTAGCAACTGCAACCGGTGTACATTCTCAGGTCCAACTGCAGCAGCC |
| T4hmFH_V | CTAGTAGCAACTGCAACCGGTGTACATTCTSAGGTYCAGCTGCARCAGTC |
| T4hmFH_VI | CTAGTAGCAACTGCAACCGGTGTACATTCTCAAGTGCAGATGAAGGAGTC |
| T4hmFH_VII | CTAGTAGCAACTGCAACCGGTGTACATTCTCAGATCCAGTTGGYGCAGTC |
| T4hmFH_VIII | CTAGTAGCAACTGCAACCGGTGTACATTCTCAGGTCCAACTCCAGCAGCC |
| T4hmFH_IX | CTAGTAGCAACTGCAACCGGTGTACATTCTCAGGTGCAACTGAAGCAGTC |

**Forward for mouse (for IgG1 expression vector) Specific clonotype**

| T4hmFH_10 | CTAGTAGCAACTGCAACCGGTGTACATTCTGATGTACAGCTTCAGGAGTCAGG |
| --- | --- |
| T4hmFH_11 | CTAGTAGCAACTGCAACCGGTGTACATTCTGAGGTCCAGCTGCAACAATCTGG |
| T4hmFH_12 | CTAGTAGCAACTGCAACCGGTGTACATTCTCAGGTCCAACTGAAGCAGTCTGG |
| T4hmFH_13 | CTAGTAGCAACTGCAACCGGTGTACATTCTCAGGTCCAGCTGCAGCAGTC |

**Multiplex reverse (for mouse IgG1 expression vector)**

| T4mRG_I | TGGGCCCTTGGTGGTGGCGCTCGAGACGGTGACCGTGGTCCCT |
| --- | --- |
| T4mRG_II | TGGGCCCTTGGTGGTGGCGCTCGAGACTGTGAGAGTGG**TGCCAGG** |
| T4mRG_III | TGGGCCCTTGGTGGTGGCGCTCGAGACAGTGACCAGAG**TCCCTTGG** |
| T4mRG_IV | TGGGCCCTTGGTGGTGGCGCTCGAGACGGTGACTGAGG**TTCCTTGA** |

**Multiplex forward (for mouse IgK expression vector)**

| T4hmFK_I | CTAGTAGCAACTGCAACCGGTGTACATTCCGAAAWTGTGCTCACCCAGTC |
| --- | --- |
| T4hmFK_II | CTAGTAGCAACTGCAACCGGTGTACATTCCCAAATTGTTCTCACCCAGTC |
| T4hmFK_III | CTAGTAGCAACTGCAACCGGTGTACATTCCRACATTGTGCTGACCCAATC |
| T4hmFK_IV | CTAGTAGCAACTGCAACCGGTGTACATTCCGAAACAACTGTGACCCAGTC |
| T4hmFK_V | CTAGTAGCAACTGCAACCGGTGTACATTCCGATATTGTGATGACSCAGGC |
| T4hmFK_VI | CTAGTAGCAACTGCAACCGGTGTACATTCCRRTRTTGTGATGACCCARAC |
| T4hmFK_VII | CTAGTAGCAACTGCAACCGGTGTACATTCCGATATCCAGATGACACAGAC |
| T4hmFK_VIII | CTAGTAGCAACTGCAACCGGTGTACATTCCGACATTGTGATGACMCAGTC |
| T4hmFK_IX | CTAGTAGCAACTGCAACCGGTGTACATTCCGACATCCAGATGACHCAGTC |

**Multiplex reverse (for mouse IgK expression vector)**

| T4mRK_I | GGATACAGTTGGTGCAGCATCCGTACGTTTCAGCTCCAGCTTGGTCCC |
| --- | --- |
| T4mRK_II | GGATACAGTTGGTGCAGCATCCGTACGTTTTATTTCCAGTCTGGTCCC |
| T4mRK_III | GGATACAGTTGGTGCAGCATCCGTACGTTTKATTTCCARCTTKGTSCC |
