## Supplementary Table 2 for "FB5P-seq-mAbs: monoclonal antibody production from FB5P-seq libraries for integrative single-cell analysis of B cells"

**Supplementary Table 2 : Primers for screening PCR and sequencing**

| Screening Forward Primers (IgG1 and IgK) | GCTTCGTTAGAACGCGGCTAC |
| --- | --- |
| Screening Reverse Primers (IgG1) | GGGTCACCATGGAGTTAGTTTGG |
| Screening Reverse Primers (IgK) | TCCACTTGACATTGATGTCTTTGG |
