## Supplementary File 1 for "FB5P-seq-mAbs: monoclonal antibody production from FB5P-seq libraries for integrative single-cell analysis of B cells"

**Reagent setup**

**PBS:** Dilute ten-fold the initial 10X PBS to obtain de final solution of PBS at 1X.

**OVA:** To prepare OVA from stocks (lyophilized). Equilibrate OVA at room temperature. Put at 10 mg/ml (add H_2_O nuclease free). Vortex gently until you see the solution is clear.

**OVA immunization:** For each mouse, add 10 µl of OVA at 100 µg, 50 µl of alum and 40 µl of PBS.

**Tips:** Cut tips with a blade and pipet the corresponding volume of alum many times then add it drop by drop in the 2 ml eppendorf tube. Place in the termomixer at 21°C at 800 rpm for 30 min.

**Tamoxifen:** For each mouse, add 5 mg of tamoxifen in 200 μL of arachidic acid oil. Perform two sonication of 15 sec.

**FACS Buffer:** In a stericup to sterilize the solution with vacuum pomp, add 50 ml of PBS 10X, 10 ml of FCS (2% final), 2 ml of EDTA 2 mM and 440 ml of ddH_2_O. Mix and store medium at 4–8 °C for up to 3 months.

**Triton X-100:** It is prepared by diluting a commercially available 10% v/v with PCR-grade H_2_O.

**(dT)30_Smarter primer:** The 100 µM stock is diluted to 10 µM with PCR-grade H_2_O before preparing the lysis mix.

**External RNA Controls Consortium (ERCC) spike-ins mix**: It is kept as stock aliquots of 1 µl (1 µg/µl) at -80°C. Further dilutions (0.1 µg/µl), prepared by adding 9 µl PCR-grade H_2_O to an aliquot, can be stored at -20°C and re-used for a maximum of 5 additional freeze-thaw cycles. The 0.5 pg/µl dilution used in the lysis mix is prepared *ad hoc* from the 0.1 µg/µl dilution by serial 1:100, 1:100 and 1:20 dilutions in PCR-grade H_2_O.

**dNTP mix:** dNTP mix is prepared by mixing equal amounts of dATP, dCTP, dGTP, dTTP, each at a stock concentration of 100 mM.

**Mix of i7 primers:** Pool i7_BCx at 0.5 µM with i7_primer at 10 µM.

**Tris Borate EDTA (TBE):** Dissolve 108 g of Tris base with 55 g of boric acid and 9,3 g of EDTA in 1L of ddH_2_O. Stir until it has completely dissolved. Autoclave and store the solution at room temperature for 3 years.

**Agarose gel 0.8%:** Add 2,4 g of agarose to 300 ml of TBE. Heat the mixture in microwave until boiling to dissolve the agarose. Cool it and then add 15 µl at 10 mg/ml (7,5 µg/ml) of ethidium bromide. Pour it into the gel casting tray and let it cool until the samples are deposited.

**Ampicillin:** Add 1 g of ampicillin sodium salt at 100 mg/ml to 10 ml of ddH_2_O. Fully dissolved and sterilize the solution with a steritop. Store the solution at −20 °C for 1 year.

**LB medium-ampicillin:** Suspend 20 g in 1L of ddH_2_O. Autoclave for 15 minutes at 121°C and add 100 µg/ml of ampicillin.

**Transformation and Storage Solution (TSS):** For 100 ml you need 1 g of tryptone**,** 0.5 g of yeast extract**,** 0.5 g of NaCl**,** 10 g of polyethylene glycol 3350**,** 5 ml of DMSO**,** 5 ml of MgCl_2_ (1M) in 70 ml of ddH_2_O. Stir until dissolved. Adjust the pH to 6.5 with HCl or NaOH. Adjust to 100 mL with sterile distilled water. Sterilize by filtering through 0.22-micron steritop filter. Store at 4 °C. Stable for up to 6 months.

**Competent bacteria:**

1. Using sterile technique, streak an *E. coli* host strain (e.g., JM109) from a glycerol stock onto an LB agar plate. Incubate overnight at 37 °C.
2. Isolate a single colony and inoculate 50 to 100 mL of LB medium.
3. Incubate at 37°C with shaking at 250 rpm.
4. Grow cells to an A600 of 0.4 to 0.5.
5. Sediment the cells at approximately 2500 g for 15 min at 4°C, then gently resuspended in 1/10 volume (5 to 10 mL) of ice-cold TSS and place on ice. Store at -80°C after aliquoting to desired volume.

**LB-ampicillin agar plates:** Suspend 35 g in 1 L of ddH_2_O. Heat to boiling while stirring to dissolve all ingredients completely and add 1 g of glucose. Autoclave for 15 minutes at 121°C and 100 µg/ml of ampicillin.

**Agarose gel 1.5%:** Add 4.5 g of agarose to 300 ml of TBE. Heat the mixture until dissolve the agarose. Cool it until 45 °C, and then add 15 µl of ethidium bromide at 10 mg/ml (7,5 µg/ml). Pour it into the gel casting tray and let it cool until it has solidified. Prepare the gel on the day of use.

**Polyethyleneimine (PEI):** Dissolve in a large volume (450 ml) in a beaker with ddH_2_O 500 mg of polyethyleneimine. Adjust the pH to 7.0 with 1 M HCl. Adjust the volume of solution with ddH_2_O to the final concentration of 1 mg/mL PEI. Filter the PEI solution through a 0.22-micron steritop filters. Make 2 mL aliquots of PEI solution and store at − 80°C.

**Glucose 0.4%:** Add 500 µl to 50 ml of ddH_2_O. Stir until it has completely dissolved. Store medium at 4–8 °C for up to 3 months.

**Tryptone 0.5%:** Add 2.5 ml to 50 ml of ddH_2_O. Stir until it has completely dissolved. Store medium at 4–8 °C for up to 3 months.

**Valproic acid 0.5 mM:** Add 500 µl to 50 ml of ddH_2_O. Stir until it has completely dissolved. Store medium at 4–8 °C for up to 3 months.

**ELISA washing buffer:** Add 500 µl of Tween 20 (0.05%) in 1L of PBS. Store buffer at room temperature for up to 5 years.

**ELISA blocking buffer:** Add 4 g of BSA in 200 ml of PBS. Store blocking solution at 4–8 °C for a maximum of 2 d.

**Expi293^TM^ medium:** Add 2 ml of penicillin–streptomycin (10,000 U/ml) to 1 L of Expi293^TM^ medium. Store medium at 4–8 °C for up to 3 months.

**Glycine buffer:** Add 0.75 g of glycine to 80 ml of ddH_2_O. Stir until it has completely dissolved. Adjust the pH to 2.9 with HCl. Bring the final volume to 100 ml with ddH_2_O. Autoclave the solution and store it at room temperature for 3 years.

**Tris buffer:** Add 1.21 g of Tris base to 8 ml of ddH_2_O. Stir until it has completely dissolved. Adjust the pH to 8.0 with HCl. Bring the final volume to 10 ml with ddH_2_O. Autoclave and store the solution at room temperature for 3 years.

**TMB solution:** Mix ½ TMB + ½ Peroxyde solution (H_2_O_2_), mix and directly add to wells.
