## Supplementary File 2 for "FB5P-seq-mAbs: monoclonal antibody production from FB5P-seq libraries for integrative single-cell analysis of B cells"

**TIMING**

A. **Immunization and treatments – Timing 2h30**

Immunization, Steps 1-3: Timing 1h00

Tamoxifen administration, Step 4: Timing 1h30

B. **Staining and cell sorting – Timing 5h00**

Process organs and cell suspensions preparation, Step 5-12: Timing 1h30

Staining, Steps 13-24: Timing 1h30

FACS sort, Steps 25-33: Timing 2h00

C. **Reverse transcription and template switching – Timing 4h00**

Prepare 2X RT mastermix containing individual barcoded TSO, Steps 34-40: Timing 30 min

Reverse transcription, Steps 41-57: Timing 3h30 min

D. **cDNA amplification by PCR – Timing 3h30**

PCR amplification of full lenght cDNA transcripts, Steps 58-65: Timing 3h30

E. **cDNA pool - Timing 30 min**

Step 66-69

F. **Purification – 1h30**

Cleanup of select cDNA products, Step 70-82: Timing 1h00

Second cleanup of select cDNA products, Step 83-94: Timing 30 min

G. **FB5P-seq library preparation – Timing 3h45**

Quantification with Qubit and BioAnalyzer, Steps 95-96: Timing 1h00

Tagmentation, Steps 97-107: Timing 45 min

Sample index PCR, Steps 108-115: Timing 1h00

PCR cleanup, Steps 116-132 – Timing 1h00

H. **Illumina sequencing – Timing ~24h00**
I. **Analysis single cell data – Timing days-month**
J. **Preparation of expression vectors for cloning – Timing 8h00**

Steps 134-147

K. **PCR amplification of the heavy and light chain genes – Timing 5h00**

Steps 148-160

L. **Cloning by SLIC of heavy and light chain genes in expression vectors – Timing 1 day**

Steps 161-182

M. **Colony screening – 1 day**

Steps 183-195

N. **DNA preparation – 1 day**

Steps 196-199

O. **Cell culture and transfection – Timing 7 days**

Steps 200-207

P. **Optional: ELISA control of production – Timing 5h00**

Steps 208-226

Q. **Recovery of culture supernatant and preparation of 50% protein G beads – Timing 1h00**

Steps 227-236

R. **Antibodies purification – Timing 4h00**

Steps 237-247

S. **Antibodies concentration – Timing 1h00**

Steps 248-253

**Step by Step procedure**

A. **Mice immunization and treatments -** **Timing 2h30**

For this experiment 8 mice were used, 2 mice for time point. The aim is to analyze GC B cell responses at J5, J10, J15 and J20 with mice at least 7 weeks of age. This experiment can be adapted.

**Immunization -** **Timing 1h00**

Immunization is performed with a 0.3 ml syringe to inject OVA first for J20 mouse, then 10 days later for J10 mouse and 5 days later for J5 mouse with OVA immunization (see “Reagent Setup”).

**Tips:** As for some syringe there is a dead volume, prepare solution for 1 more mouse.

1. Prepare OVA immunization (see “Reagent Setup”).
2. Load 100 µl into syringe.
3. Immunize on each side of tail with 50 µl.

**Tips:** The bevel of the syringe have to be in the bottom and the orifice of the syringe on the top.

**Tamoxifen administration -** **Timing 1h30**

4. Perform gavage 48h before sacrifice (see “Experimental Setup) with 200 µl of tamoxifen (see “Reagent Setup”) per mouse.

**Tips:** To reuse the gavage syringe, at the end, wash 5 times the gavage syringe with PBS. Then, wash with ethanol and put for storage in a 50 ml tube containing ethanol.

B. **Staining and cell sorting -** **Timing 5h00**

**Process organs and cell suspensions preparation -** **Timing 1h30**

5. Sacrifice mice according to all relevant governmental and institutional guidelines and regulation, in compliance with protocols approved by local animal ethics.

**Tips:** The day of sacrifice is the same for all mice to avoid batch effects. The only variation is the day of immunization.

6. Isolate dLN (inguinal and periaortic) and place in 5 ml FACS tube containing cold FACS buffer (see “Reagent Setup”).
7. Transfer LN sample into a small Petri Dish and strain into 70 µm cell strainer with syringe plunger in a Petri Dish and transfer into a 15 ml tubes.
8. Centrifuge at 300 g for 5 min at 4°C.
9. Discard supernatant and resuspend in 1 ml FACS Buffer.

**Tips:** Count cells with Trypan Blue: 190 µl of Trypan Blue and 10 µl of cells. Dispense 10 µl in the grid for counting.

10. Add 9 ml of FACS Buffer.
11. Centrifuge at 300g for 5 min at 4°C.
12. Resuspend cells in the corresponding FACS Buffer volume to have 100 x 10^6^ cells/ml.

**Staining - Timing 1h30**

Steps 13 to 24 are performed into sterile hood.

13. Prepare antibody mixes and compensation mix in FACS Buffer.

**Tips:** It’s important to use negative control and titrate each antibody to optimize the staining with labeled antibodies. To avoid background-staining use appropriate species and specific isotype control antibodies.

| Antibody Table | Clone | Final Dilution | 2X concentration |
| --- | --- | --- | --- |
| CXCR4-PerCP-eF710 | 2B11 | 1/50 | 25 |
| GL7-BV421 | GL7 | 1/500 | 250 |
| CD138-BV711 | 281-2 | 1/50 | 25 |
| OVA-AF 647 | ??? | 1/800 | 400 |
| CD3-APC-Cy7 | 17A2 | 1/100 | 50 |
| Gr1-APC-Cy7 | RB6-8C5 | 1/100 | 50 |
| CD38-PE | 90 | 1/500 | 250 |
| CD19-PE-Dazzle594 | 6D5 | 1/100 | 50 |

Composition of antibody mix :

14. Dispense cells in 5 ml FACS tubes for staining.
15. Add 1:50 (v:v) Mouse Fc block at 0.5 mg/ml to sample.
16. Protect from light and incubate for 10 min at 4°C.
17. Incubate cells with 2X antibody mix (see “Antibody Table”) or compensation mixes for 30 min at 4°C.
18. Wash with 1 ml of FACS Buffer and centrifuge at 300g for 3 min at 4°C.
19. Wash with 1 ml of PBS and centrifuge at 300g for 3 min at 4°C.
20. Add 1X Live/Dead mix.
21. Protect from light and incubate for 10 min at 4°C.
22. Wash with 1 ml of FACS Buffer and centrifuge at 300g for 3 min at 4°C.
23. Resuspend in 400 µl of FACS Buffer.
24. Filter the cells in a new 5 ml FACS tubes with pre-separation filter.

**Influx sort -** **Timing 2h00**

Steps 25 to 26 are performed into sterile hood.

25. Prepare enough LYSIS mix for 110 wells if planning to sort a full 96-well plate. For this experiment, we prepared for 20 plates and stored at -80°C.

| **LYSIS mix** | **Stock conc.** | **Final conc.** | **Volume per well (µl)** | **Wells** | **Total (µl)** |
| --- | --- | --- | --- | --- | --- |
| **Triton X-100** | 0.40% v/v | 0.10 | 0.5 | 2200 | 1100 |
| **Rnase OUT** | 40 U/µl | 1 | 0.05 | 2200 | 110 |
| **dNTP mix** | 25 mM | 1 | 0.08 | 2200 | 176 |
| **(dT)30_Smarter** | 10 µM | 2.5 | 0.5 | 2200 | 1100 |
| **ERCC spike-in Mix*** | 0.5 pg/µl | 0.0125 | 0.05 | 2200 | 110 |
| **PCR-grade H_2_O** | NA | NA | 0.82 | 2200 | 1804 |
|  |  |  | 2 |  | 4400 |

Composition of LYSIS mix (see “Reagent setup”):

* The amount of ERCC spike-in mix added to each well (0.025 pg) has been optimized for scRNAseq analysis of *ex-vivo* mouse and human B and T lymphocytes, which contain little mRNA. It may need to be adjusted when analyzing cells with more mRNA.

26. Dispense **2 µl** of LYSIS mix into MicroAmp EnduraPlate Optical 96-Well plates in each well and store at -80°C until several months.

**Tips:** Prepare in advance enough plaque containing lysis mix for cell sorting. You should keep them in -80°C in aluminum paper up to months.

27. Thaw lysis mix plates for sorting.
28. The sorting is performed in BD Influx^TM^ cell sorter.

**Tips:** These gates should be adjusted depending of your experiments.

29. Sort 1 cell per well by BD Influx^TM^ cell sorter into 96-well PCR plate.

**Notes:** For this experiment, we sorted 18 plates.

30. After sorting, cover sorting plates with adhesive film.
31. Briefly centrifuge the plate.
32. Immediately freeze plates in dry ice.
33. Transfer to -80°C freezer.

***Pause Point:*** *After the collection of cells, immediately freeze plates in dry ice. Plates can be stored several months at -80°C.*

C. **Reverse transcription and template switching - Timing 4h00**

**Tips:** To prevent any contamination, all steps are performed into single-cell hood. This mean a designated hood as DNA/RNA-free to prepare solution for reverse transcription and PCR reactions mixes. The hood, all surfaces and all reagents must be cleaned with 10% of bleach. Use sterile filter pipette tips.

The reverse transcription (RT) step uses 96 different well-specific template-switching oligonucleotides (TSO) to introduce a well-specific DNA barcode in the 5’-end of cDNAs.

The 2X RT mastermix 96-well plate can be prepared in advance with enough reagents for processing several plates. It stored at -20°C and can be re-used for maximum of 5 freeze-thaw cycles.

**Prepare 2X RT mastermix containing individual barcoded TSO – Timing 30 min**

34. Prepare enough mastermix for 20 plates. Freeze and re-use max 5 times.

**Tips:** SuperScript II / RNase OUT is added at step 50 after primer annealing.

Composition of 2X RT mastermix:

| **2X RT mastermix** | **Stock conc.** | **Final conc.** | **Volume per well (µl)** | **Runs** | **Total (µl)** | **Wells** | **Premix (µl)** |
| --- | --- | --- | --- | --- | --- | --- | --- |
| **SuperScript II Buffer** | 5 X | 2 | 1 | 20 | 20 | 110 | 2200 |
| **DTT** | 100 mM | 10 | 0.25 | 20 | 5 | 110 | 550 |
| **Betaine** | 5 M | 2 | 1 | 20 | 20 | 110 | 2200 |
| **MgCl_2_** | 1000 mM | 12 | 0.03 | 20 | 0.6 | 110 | 66 |
| **TSO_BCx_UMI5_TATA_PM*** | 100 µM | 5 | 0.13 | 20 | 2.5 |  |  |
| **PCR-grade H_2_O** |  |  | 0.1 | 20 | 1.9 | 110 | 209 |
|  |  |  | 2.5 |  | 50 |  | 5225 |

* TSO_BCx_UMI5_TATA_PM are DNA-RNA oligonucleotides each containing a 5’-end 21-nucleotide universal PCR handle, followed by an 8-nucleotide long well-specific barcode, a 5-nucleotide long random Unique Molecular Identifier (UMI), a TATA insulator, and terminated by 3 riboguanine (rG) bases at the 3’-end.

35. Prepare 2X RT mastermix plate into MicroAmp EnduraPlate Optical 96-Well plates.
36. Dispense **47.5 µl** per well of premix with 8-micropipette 100 µl.
37. Add **2.5 µl** of 100 µM TSO (stock) with 8-channel micropipette 10 µl.
38. Cover sorting plates with adhesive film compatible with PCR.
39. Briefly centrifuge the plate.
40. Store at -20°C.

**Reverse transcription – Timing 3h30**

41. Thaw 2 plates containing single cells in lysis mix from -80°C into ice.

**Tips:** You should do minimum 2 plates to not pipette too small volumes.

42. Briefly centrifuge the plate.

**Tips:** Heat the lid before at 72°C.

43. Place in Thermal Cycler with following program:

| PCR machine |  |
| --- | --- |
| 3 min at 72°C | *This step hybridizes the (dT)30_Smarter primer* |

44. Immediately place the plates back on ice until step 52.
45. Briefly spun down in a benchtop plate centrifuge.
46. Prepare RT mix plate sufficient for 2 plates on ice.

Composition of 1X RT mastermix:

| 1X RT mastermix | | Stock Conc. | Final Conc. | Volume per well (µl) | # wells | Total (µl) | #plates | Total (µl) |
| --- | --- | --- | --- | --- | --- | --- | --- | --- |
| SuperScript II (SSII) RT | 200 U/µl | | 10 | 0.25 | 220 | 55 |  | 0 |
| Rnase OUT | 40 U/µl | | 2 | 0.25 | 220 | 55 |  | 0 |
| 2X RT mastermix* | 2 | | 1 | 2.5 |  |  | 2 | 5 |
| Lysis Product ** |  | |  | 2 |  |  |  |  |
| Total |  | |  | 5 |  | 110 |  | 5 |

***** Pipeted from the 2X RT mastermix plate previously thawed on ice into the corresponding well of the new plate. New tips should be used for each well to avoid barcode cross-contamination.

** The SSII/Rnase OUT will be added to the lysis product to avoid material loss.

47. Thaw on ice 2X RT mastermix plate.
48. Pool in a tube the SSII and Rnase OUT and dispense 13 µl of mix into 8-tube PCR strip.
49. In a new plate (1X RT mastermix) add **5 µl** of 2X RT mastermix with 8-channel micropipette 10 µl for 2 plates.
50. Add in the 1X RT mastermix plate **1 µl** of SSII/Rnase OUT with 8-channel micropipette 10 µl for 2 plates. Cover plates with adhesive film.

**Tips:** Change tips to avoid contamination.

51. Vortex and briefly centrifuge plate.

Dispense RT mastermix into sample plates :

52. After removing the adhesive cover, using 8-channel micropipette 10 µl, dispense **3 µl** of RT mix into plates containing single cells in lysis from step 44.

**Tips:** Change tips to avoid contamination.

53. Cover plate with adhesive film compatible with PCR.
54. Vortex and briefly spun down in a benchtop plate centrifuge.

**Tips:** Heat the lid before at 70°C.

55. Place in Thermal Cycler with following program:

| PCR machine |  |
| --- | --- |
| 90 min at 42°C | *RT and template switching* |
| For 10 cycles  2 min at 50°C  2 min at 42°C | *Unfolding of RNA secondary structures*  *Completion/continuation of RT and template switching* |
| 15 min at 70°C | *Enzyme inactivation* |
| Hold at 4°C | *Safe storage* |

56. Briefly centrifuge the plate.
57. Perform long distance PCR (LD-PCR) amplification of full transcripts or store at -20°C for few days.
D. **cDNA amplification by PCR -** **Timing 3h30**

**LD-PCR amplification of full transcripts**

58. Prepare 7.5 µl of PCR mix per sample.

Composition of LD-PCR Mix :

| **LD-PCR Mix** | **Stock Conc.** | **Final Conc.** | **Volume per well (µl)** |  | **Wells** | **Total (µl)** |
| --- | --- | --- | --- | --- | --- | --- |
| **KAPA HiFi ReadyMix** | 2 X | 1 | 6.25 |  | 220 | 1375 |
| **PCR_Satija*** | 20 µM | 0.2 | 0.13 |  | 220 | 27.5 |
| **SmarterR*** | 20 µM | 0.2 | 0.13 |  | 220 | 27.5 |
| **Lysis + RT-TS **** |  |  | 5 |  |  |  |
| **PCR grade H2O** |  |  | 1 |  | 220 | 220 |
|  |  |  | 12.5 |  |  | 1650 |

* LD-PCR primers should be order with HPLC.

** The LD-PCR mix will be added to the plate containing the cDNA products to avoid material loss.

59. Aliquot LD-PCR mix into an 8-tube strip, 206 µl per tube.
60. Dispense **7.5 µl** PCR mix into LYSIS+RT-TS samples for 2 plates with 8-channel 10 µl micropipette. Keep on ice in PCR cooling block.

**Tips:** Change tips to avoid contamination.

61. Cover plate with adhesive film compatible with PCR.
62. Vortex and briefly centrifuge plate.

**Tips:** Heat the lid before at 98°C.

63. Place in Thermal Cycler with following program:

PCR machine

3 min at 98°C

For 22 cycles

15 sec at 98°C

20 sec at 67°C

6 min at 72°C

5 min at 72°C

Hold at 4°C

**Tips:** The number of PCR cycles (22) has been optimized for scRNAseq of *ex-vivo* mouse and human B and T lymphocytes, which contain little mRNA. It may need to be adjusted when analyzing cells with more mRNA.

64. Briefly centrifuge plate.
65. Store at -20°C until CleanNGS Beads purification or BCR amplification for antibodies production.

**Tips:** You should do all plates until this step to perform library at the same time.

***Pause point:*** *After LD-PCR plates can be stored several years at -20°C.*

E. **cDNA pool -** **Timing 30 min**
66. Take **5 µl** from each well of the plate and for each plate put them in a tube DNA LoBind 1.5 ml.

**Note:** We have 18 plates to pool. We obtained one tube per plate.

67. With the p1000 assess the total volume obtained for each tube and complete to **500 µl** with PCR-grade H_2_O.
68. Prepare a strip of PCR tubes and purify 100 µl of full-length cDNA products.
69. Keep the remaining 400 µl in a tube DNA LoBind 1.5 ml and store at -20°C.

***Pause point:*** *After cDNA pool, tubes can be stored several weeks at -20°C.*

F. **Purification -** **Timing 1h30**

**CleanNGS Beads cleanup of select cDNA products -** **Timing 1h00**

**Note:** Purify full length cDNA products to remove primer dimers.

70. Equilibrate CleanNGS Beads at RT for 30 min before starting.
71. Add **60 µl** (0.6X) of CleanNGS Beads in each PCR tube containing 100 µl of full-length cDNA.
72. Pipette up and down 10 times to mix with a p200 adjusted to 150.
73. Incubate 5 min at RT.
74. Place on magnet during 5 min to collect beads on side of the tube.
75. Discard beads supernatant and wash beads on magnet twice with **200 µl** freshly prepared 80% ethanol in PCR grade H_2_O.

**Tips:** Be careful while putting ethanol to not remove the beads from the side of the tube.

76. Centrifuge tubes to remove ethanol.
77. Remove again the remaining ethanol with a p10 to increase the yield (on magnet).
78. Dry beads for maximum 10 minutes or until cracks appear.
79. Resuspend well beads in **100.5 µl** PCR-grade H_2_O.
80. Incubate for 5 min.
81. Place on magnet during 2 min to collect beads on side of the tube.
82. Aspirate **100 µl** purified cDNA and collect in a new PCR tube.

**CleanNGS Beads second cleanup of select cDNA products -** **Timing 30 min**

83. Add **60 µl** (0.6X) CleanNGS Beads in each tube containing 100 µl of purified cDNA.
84. Pipette up and down 10 times to mix with a p200 adjusted to 150.
85. Incubate for 5 min at RT.
86. Place on magnet during 5 min to collect beads on side of the tube.
87. Remove bead supernatant and wash beads on magnet twice with 200 µl freshly prepared 80% ethanol in PCR grade H_2_O.

**Tips**: Be careful while putting ethanol to not remove the beads from the side of the tube.

88. Centrifuge tubes to remove ethanol.
89. Remove again the remaining ethanol with a p10 to increase the yield (on magnet).
90. Dry beads for 10 minutes or until cracks appear.
91. Resuspend well beads in **15.5 µl** PCR-grade H_2_O.
92. Incubate for 5 min.
93. Place on magnet during 2 min to collect beads on side of the tube.
94. Aspirate **15 µl** purified cDNA and collect in a new PCR tube.

***Pause point:*** *After purification tubes can be stored up to several months at -20°C.*

G. **FB5P-seq library preparation - Timing 3h45**

For 5’-end sequencing library preparation, 800 pg purified cDNA pool are processed with the Nextera XT DNA sample Preparation kit (Illumina FC-131-1096), according to the manufacturer’s instructions with modifications.

95. Quantify with Qubit and BioAnalyzer High Sensitivity DNA test.
96. For the library, prepare an aliquot of cDNA for each tube at 0.2 ng/µl with PCR-grade H_2_O.

**Tagmentation - Timing 45 min**

97. Thaw ATM, TD and input DNA on ice.
98. Remove the NT buffer from the fridge and set it to RT.
99. Prepare tagmentation mix for 18 tubes.

Composition of tagmentation mix:

| Tagmentation mix | Volume (µl) | | Wells | Total (µl) |
| --- | --- | --- | --- | --- |
| TD buffer | 10 | 18.5 | | 185 |
| ATM | 5 | 18.5 | | 92.50 |
| Total | 15 |  | | 278 |

100. In a new PCR strips, add **4 µl** of diluted cDNA at 0.2 ng/µl (800 pg) of sample DNA to each tube.
101. Add **1 µl** of PCR grade H_2_O to each tube to have 5 µl of final volume.
102. Add **15 µl** of tagmentation mix to each tube.
103. Pipette up and down 5 times to mix and centrifuge.

**Tips:** Change tips between samples.

**Tips:** Heat the lid before at 55°C.

104. Place in Thermal Cycler with following program:

PCR machine

10 min at 55°C

Hold at 10°C

105. When temperature reaches 10°C, remove tubes from the PCR machine.
106. Proceed immediately to neutralization by adding **5 µl** of NT buffer. Pipette up and down 5 times to mix.

**Tips:** Change tips between samples.

107. Centrifuge and incubate 5 min at RT.

**Sample index PCR -** **Timing 1h00**

Sample index PCR is used to enrich 5’-ends of tagmented cDNA and append barcoded Illumina sequencing adapters with the Nextera i5 primer S5xx (Illumina FC-131-2001 or FC-131-2002) and i7 primers mix (i7_BCx at 0.5 µM + i7_primer at 10 µM). When performing parallel analysis on several 96-well plates of single cells, the pooled cDNA from each plate is amplified with a distinct i7_BCx primer.

108. Thaw index primers at RT.
109. Thaw NPM on ice.
110. Add **5 µl** of index primer i5 mix in each tube.
111. Add **5 µl** of index primer i7 mix in each tube.
112. Add **15 µl** of NPM mix in each tube.

**Tips:** Change tips between tubes.

113. Pipette up and down 5 times to mix.
114. Centrifuge.

**Tips:** Heat the lid before at 95°C.

115. Place in Thermal Cycler with following program:

PCR machine

3 min at 72°C

30s at 95°C

For 12 cycles

10 sec at 95°C

30 sec at 55°C

30 sec at 72°C

5 min at 72°C

Hold at 4°C

**CleanNGS Beads cleanup -** **Timing 1h00**

116. Thaw beads and RB illumina.
117. The final volume of PCR is 50 µl, add **50 µl** of water to complete to 100 µl for purification.
118. Add 80 µl (0.8X) CleanNGS Beads in each tube containing 100 µl of full-length cDNA.
119. Pipette up and down 10 times to mix with a p200 adjusted to 150.
120. Incubate 5 min at RT.
121. Place on magnet during 5 min to collect beads on side of the tube.
122. Remove bead supernatant and wash beads on magnet twice with 200 µl freshly prepared 80% ethanol in PCR grade H_2_O.

**Tips:** Be careful while putting ethanol to not remove the beads from the side of the tube.

123. Centrifuge tubes to remove ethanol.
124. Remove again the remaining ethanol with a p10 to increase the yield (on magnet).
125. Dry beads for 10 minutes or until cracks appear.
126. Resuspend well beads in **20.5 µl** RB illumina.
127. Incubate for 5 min.
128. Place on magnet during 2 min to collect beads on side of the tube.
129. Aspirate **20 µl** purified cDNA and collect in a new PCR tube.
130. Quantify with Qubit HS DNA test.
131. Store the strip at -20°C until Bioanalyzer High Sensitivity DNA chip analysis.

*See troubleshooting*

**Tips:** The resulting library is expected to have a broad size distribution (300-1000 bp) with an average size of 600-800 bp when controlled on a BioAnalyzer with High Sensitivity DNA Analysis Kits.

132. Prepare pool following platform recommandation.
H. **Illumina sequencing – Timing 1 day**

Libraries generated from multiple 96-well plates of single cells and carrying distinct i7 barcodes may be pooled for sequencing. For optimal results, 5x10^5^ reads per cell should be targeted in custom paired-end single-index Illumina sequencing with the following primers and cycles: read 1 (Read1_SP_PM, 67 cycles), read i7 (i7_SP_PM, 8 cycles), read 2 (Read2_SP_PM, 16 cycles).

I. **Analysis single cell data- Timing days-month**
133. [GitHub - MilpiedLab/FB5P-seq](https://github.com/MilpiedLab/FB5P-seq)
J. **Preparation of expression vectors for cloning - Timing 8h00**

For heavy and light chain cloning, the heavy chain IgG1 mouse expression vector and the light chain IgK mouse expression vectors were provided by the Nussenzweig laboratory (Von Boehmer et al, 2016). Users recommended to digest vectors in advance.

**Tips:** Handle restriction enzyme on ice.

**For mouse IgG1 vector:**

**Tips:** Both vectors used are at 1 µg/µl.

134. Digest **10 µl** of mouse IgG1 vector with **5 µl** of AgeI-HF and **5 µl** of XhoI in the presence of **10 µl** of CutSmart buffer and **80 µl** of PCR-grade H_2_O at 37° for at least 3h.
135. To inactivate the enzyme, incubate the product of digestion for 20 min at 65C°.
136. Add **0.5 µl** of alkaline phosphatase and incubate at 37°C for 30 to 60 min.

**For mouse IgK vector:**

137. Digest **10 µl** of mouse IgK vector with **5 µl** of Age1-HF in the presence of **10 µl** of CutSmart buffer and **85 µl** of PCR-grade H2O at 37°C for at least 3h.
138. To inactivate the enzyme, incubate the product of digestion for 20 min at 65°C.
139. Cleanup the vector on column with appropriate kit such as QIAquick Gel Extration kit.
140. Digest the elution with **5 µl** of BsiWI in the presence of **10 µl** of NEBuffer 3.1 and add PCR-grade H_2_O until 100 µl for 3h at 55°C.
141. To inactivate the enzyme, incubate the product of digestion at 65°C for 20 min.
142. Add **0.5 µl** of alkaline phosphatase and incubate at 37°C for 30 to 60 min.

**For both vectors:**

143. Prepare a 0.8% agarose gel (see “Reagent Setup”).
144. Load the two digested vectors and migrate at 120 volts in TBE for 30 min (see “Reagent Setup”).
145. Expose the gel to UV light and cut the appropriate bands of linearized vectors and purify them with the QIAquick Gel Purification Kit.

**Tips:** Use a new scapel for different vectors to avoid contamination. Expose your vector to UV light short as possible to avoid DNA damage.

146. Measure the concentration of each linearized vectors by optical density with Nanodrop at 260 nm or equivalent.
147. Store vectors at -20°C.

***Pause point:*** *“Restricted” vectors can be stored until 3 years at -20°C.*

K. **PCR amplification of the heavy and light chain genes - Timing 5h00**

The amplification of the light and heavy chains (48 each) of antibodies are amplified in a one round of PCR.

Here, we chose to use this type of multiplex PCR Van Boehmer paper (). despite the fact that the BCR are already sequenced, in order to reduce the time and cost compared to specific sequences primer for each chain.

**Tips:** From this step, all PCR mixes are done in a DNA/RNAse free hood. All pipettes are dedicated to the preparation of PCR mix. Use filter tips.

148. Recover the cDNA from the - 20°C (from step 65) or - 80°C for long term storage and thaw on ice.
149. Take **2 µl** of the amplified cDNA and put them in a new 96 wells PCR plate with 18 µl nuclease-free water.

**Tips:** For this production, we take 48 cDNA.

150. Prepare PCR 2 mixes one for heavy chains and one for light chains.

Composition of PCR mix :

| **PCR Mix** | **Stock Conc.** | | **Final Conc.** | **Volume per well** | | **Wells** | | **Total (µl)** | |
| --- | --- | --- | --- | --- | --- | --- | --- | --- | --- |
| **PCR-grade H_2_O** |  |  | | | 30.65 | | 52 | | 1593 |
| **5X Buffer** | 5 | 1 | | | 10 | | 52 | | 520 |
| **MgCl_2_ solution** | 25 | 8 | | | 4 | | 52 | | 208 |
| **dNTPs (25mM)** | 25 | 0.3 | | | 0.6 | | 52 | | 31.2 |
| **Multiplex Forward Primers** | 50 | 0.3 | | | 0.3 | | 52 | | 15.6 |
| **Multiplex Reverse Primers** | 50 | 0.2 | | | 0.2 | | 52 | | 10.4 |
| **Go-Taq G2 Flexi enzyme** | 5 | 1.25 | | | 0.25 | | 52 | | 13 |
| **cDNA*** |  |  | | | 4 | |  | |  |
| **Total** |  |  | | | 50 | |  | | 2392 |

*cDNA is directly added in PCR plates after PCR mix.

151. Dispense **46 µl** per well of PCR mix with a multichannel pipette.
152. From each diluted well, take **4 µl** of cDNA (for heavy chain) and add them from 1 to 48 wells.
153. From each diluted well, take **4 µl** of the same cDNA (for light chain) and place them from 49 to 96.
154. Cover plate with adhesive film compatible with PCR.
155. Vortex and briefly spun down in a benchtop plate centrifuge.

**Tips:** Heat the lid before at 95°C.

156. Place in Thermal Cycler with following program:

PCR machine

3 min at 95°C

For 40 cycles

20 sec at 95°C

30 sec at 54°C

55 sec at 72°C

10 min at 72°C

Hold at 16°C

**Tips:** We added nucleotides to the VH reverse primers to perform only one PCR for IgH and IgK with the same temperature.

157. Prepare a 2% agarose gel (see “Reagent Setup”).
158. Load **5 µl** from each well on gel with the 1Kb Plus DNA ladder on each row and migrate at 120 volts in TBE for 30 min (see “Reagent Setup”). During this time, keep the plate with adhesive film and place it at 4°C.

*See troubleshooting*

**Tips:** The Kappa chain is approximately 380 bp and 430 bp for heavy chain.

**Tips:** Some IgH sequences from the same clonotype are not recognized by multiplex primers. For these sequences we designed specific primers.

159. Purify the amplified chain using a commercial 96-well kit such as QIAquick 96 PCR purification kit and elute the material into **40 µl**.
160. Measure the concentration of each amplified chain by optical density with Nanodrop at 260 nm or equivalent.

**Tips:** At this stage the concentration of the fragments should be adjusted to 40 ng/µl.

***Pause point:*** *After the quantification of amplified chain, plates can be stored several years at -80°C.*

L. **Cloning by SLIC of heavy and light chain genes in expression vectors. Timing 24h00**

By using previously prepared expression vectors, the SLIC cloning method allows the assembly of inserts in vectors to be performed quickly and easily in a single homologous recombination reaction.

161. Thaw IgG1 and IgK vectors at room temperature.
162. Place a 96-well PCR plate on ice for at least 10 minutes beforehand.
163. Adjust concentration of purified DNA to 40 ng/µl with nuclease-free water.
164. Prepare a plate with **1 µl** of purified DNA at 40 ng/µl.

**Tips:** A vector to PCR product ratio of 1:5 is generally a good ratio for SLIC.

165. Thaw on ice the chemically competent E.Coli JM109 (see “Reagent Setup”) cells from -80°C and thaw on ice.
166. Prepare the SLIC mix reaction on ice.

Composition of SLIC mix:

| **SLIC Mix** | **Stock Concentration** | **Final Concentration** | **Volume per well** | **Wells** | **Total (µl)** |
| --- | --- | --- | --- | --- | --- |
| **Nuclease-free water** |  |  | 6.8 | 105 | 714 |
| **Linearized vector (40ng/µl)** | 40 | 4 | 1 | 105 | 105 |
| **NEB Buffer 2 (10X)** | 10 | 1 | 1 | 105 | 105 |
| **T4 DNA Polymerase (3000U/ml)** | 3000 | 15 | 0.2 | 105 | 21 |
| **Purified PCR*** |  |  | 1 |  |  |
| **Total** |  |  | 10 |  | 945 |

167. Add **8 µl** of the SLIC Mix with the appropriate vector to the plate with the inserts with a multichannel pipette in ice.

**Tips:** Heat the lid before at 24°C.

168. Immediately incubate the plate in the thermal cycler at 24°C for 2.5 min.
169. Immediately return to the ice for at least 10 minutes.

**Tips:** Heat the lid before at 42°C for the transformation stage.

170. Add **40 µl** of chemically competent E.Coli JM109 bacteria in new 96-well PCR plate.
171. Add **4 µl** of SLIC reaction to the competent cells with a multichannel pipette.
172. Cover plate with adhesive film.
173. Vortex and briefly spun down in a benchtop plate centrifuge.
174. Incubate the plate on ice for 30 min.
175. Put the plate on thermal cycler at 42°C for 45 sec.
176. Incubate the plate in ice for 2 min.
177. Add **50 µl** of SOC medium for each well with a multichannel and cover with adhesive film.
178. Cover plate with adhesive film.
179. Place the plate in a shaker at 37°C with a 45° angle for a better agitation. Shake for 45 to 60 min.

**Tips:** Label prepared LB-ampicillin agar plates (see “Reagent Setup) with the well number and prewarm at 37°C.

180. After incubation. briefly centrifuge the transformation plate.
181. Collect the appropriate number of LB-ampicillin agar plates and put each well in the corresponding number of LB-ampicillin agar plates.
182. Incubate LB-ampicillin agar plate at 37°C for 16h.

*See troubleshooting*

***Pause point:*** *The* *LB-ampicillin agar plates can be sealed with parafilm and stored at 4°C for 1 month.*

M. **Colony screening - Timing 24h00**

To verify the integrity of the heavy and light chains and that the sequences match those selected a PCR screen is performed on the colony. Forward and reverse primers are used (see Table 4). The primers are located upstream to the cloned variable chains and inside the constant region respectively.

183. Prepare the screening PCR Mix on ice for 2 plates.

Composition of the screening PCR Mix for 1 plate:

| **Screening PCR Mix** | **Stock Conc.** | **Final Conc.** | **Volume per well** | **Wells** | **Total (µl)** |
| --- | --- | --- | --- | --- | --- |
| **PCR-grade H_2_O** |  |  | 32.75 | 105 | 3438.75 |
| **Buffer** | 5 X | 1 | 10 | 105 | 1050 |
| **MgCl2 solution** | 25 mM | 8 | 4 | 105 | 420 |
| **dNTPs** | 25 mM | 0.5 | 1 | 105 | 105 |
| **Screening Forward Primers** | 10 mM | 0.2 | 1 | 105 | 105 |
| **Screening Reverse Primers*** | 10 mM | 0.2 | 1 | 105 | 105 |
| **Go-Taq G2 Flexi enzyme** | 5U/µl | 1.25 | 0.25 | 105 | 26.25 |
| **Total** |  |  | 50 |  | 5250 |

* Screening reverse primers are different for IgH and IgK.

184. Add **50 µl** of screening PCR mix in a PCR plate on ice.
185. Pick up a bacterial colony with the tip of a pipette of the first labeled LB-ampicillin agar plates and rotate the tip in the PCR mix well.
186. Use the same pipette tip to make an imprint on the LB-ampicillin agar plates grid with numbers for each colony.
187. Repeat these two steps for all LB-ampicillin agar plates.

**Tips:** For each agar plate,2 colonies from each LB-ampicillin agar plates are examined by colony PCR to see if they have the correct variable region.

188. Incubate the LB-ampicillin agar plates grid at 37°C for 16h.
189. Cover plate with adhesive film compatible with PCR.
190. Vortex and briefly spun down in a benchtop plate centrifuge.

**Tips:** Heat the lid before at 95°C.

191. Place in Thermal Cycler with following program:

PCR machine

3 min at 95°C

30s at 95°C

For 40 cycles

30 sec at 95°C

30 sec at 55°C

30 sec at 72°C

10 min at 72°C

Hold at 16°C

192. Prepare a 2% agarose gel (see “Reagent Setup”).
193. Load **5 µl** from each well on gel with the 1Kb Plus ladder on each row and migrate at 120 volts in TBE (see “Reagent Setup”). During this time. keep the plate with adhesive film and place it at 4°C.

**Tips:** The expected bands are around 380 bp for light chains and 450 bp for heavy chains.

194. After verification, sequence by Sanger sequencing the positive bands with the 5' screening forward primers.
195. Align the variable sequence chains with the original sequences obtained by single RNA seq.

*See troubleshooting*

N. **DNA preparation – Timing 1 day**
196. For each pair of corresponding heavy and light chains, recover the bacterial colonies screened positive by PCR and sequencing on agar plates.
197. Place one bacterial colony in 50 ml of liquid LB medium-ampicillin (see “Reagent Setup”) and culture for 16h in shaker at 37°C rotating at 180 rpm.
198. Prepare the plasmid DNA by following manufacturer’s protocol of Qiagen High Speed midi kit or equivalent to have enough DNA for transfection in 50 ml of Expi 293 culture.
199. Measure the concentration of the DNA by optical density with Nanodrop at 260 nm or equivalent.

**Tips:** The typical yield is around 150-300 µg in final volume of 500 µl. This quantity is enough for several productions of 50 ml.

O. **Culture cell – Timing 7 day**

The cells used for transfection are Expi293^TM^ grown in Expi293^TM^ Medium (see “Reagent Setup”). From one flask of 50 ml of production, the typical quantity yield is around 150 to 400 µg (3 to 8 mg/ml) after concentration in 50 µl,

Thaw an aliquot of cells and put in culture at least 2 weeks before transfection in incubator at 37°C rotating at 150 rpm with 5% CO_2_ in humid conditions. Make a passage every two days to have 0.3 x 10^6^ cells/ml.

Steps 200 to 205 are performed in a tissue culture hood.

**Transfection – Timing 3h00**

200. The day before transfection. put in culture 50 ml of Expi293^TM^ cells at 0.8x10^6^ cells/ml in 250 ml sterile flasks with vented caps.

**Tips:** Use one flask per antibody to produce. Be careful to not mix heavy and light chain of different antibodies.

201. Place flasks in a 37°C incubator rotating at 150 rpm with 5% CO_2_ under humid conditions.
202. The day of transfection in 15 ml tubes mix 20 μg of heavy chain-containing vectors with 25 μg of light chain-containing vectors for one antibody and add PBS to a final volume of 2.3 ml. Vortex the mixture for 15 seconds.
203. Add 170 μl of polyethyleneimine (PEI) at 1 mg/ml into the previous mixture for each tube. The final volume is 2.3 ml. Vortex for 15 sec.
204. Remove flasks containing the cells from the incubator and immediately add the PBS 1X-DNA-PEI mix to the flasks.
205. Mix well and return to the incubator rotating at 150 rpm at 37°C with 5% CO_2_ under humid conditions for 24 h.

**Tips:** It is important to gently rock cell culture to ensure distribution of the transfection complexes within flask.

206. The day after transfection. add 500 μl of glucose 0.4 %. 2.5 ml of tryptone 0.5 % and 500 μl of valproic acid 0.5 mM all diluted in Expi293^TM^ medium to transfected cells (see "Reagent Setup").
207. Return flasks to the incubator for another 5 days at 37°C at 150 rpm with 5% CO_2_ under humid conditions.

**Tips:** Step 208 to 226 were added to Von Boehmer protocol to save money if there were some issues with transfection which give not production of antibodies.

P. **Optional: ELISA control of production – Timing 5h00**

**Tips:** Use 8-channel micropipettes during all experiment and reservoirs for multichannel pipettes.

208. 4 days after nutrients addition. collect 500 μl of culture supernatant to test the production by ELISA.
209. Centrifuge 500 μl of culture supernatant of each flask (and untransfected cultured cells to make a negative control) at 800 g for 5 min.
210. Coat 200 μl of the culture supernatant in duplicate at the bottom of the wells of 96-well ELISA plates.
211. Cover and incubate overnight at 4°C.
212. Discard the culture supernatant by flicking the plate and tap the plate on absorbent paper to remove residual liquid.
213. Wash each well 3 times with 200 μl of ELISA washing buffer (see "Reagent Setup"). Tap the plate on absorbent paper to remove excess of PBS tween.
214. Block wells with 100 μl of blocking buffer (see "Reagent Setup").
215. Incubate for 2h at room temperature.
216. Discard the blocking buffer by flicking the plate and tap the plate on absorbent paper to remove residual liquid.
217. Wash each well 3 times with washing buffer (see "Reagent Setup"). Tap the plate on absorbent paper to remove excess of PBS tween.
218. Add HRP-coupled anti-mouse IgG secondary antibody diluted to 4000-fold in blocking buffer (see "Reagent Setup").
219. Protect from light and incubate for 2h at room temperature.
220. Equilibrate TMB solution at room temperature 20 min before adding it in the ELISA plate.
221. Discard the secondary antibody by flicking the plate and tap the plate on absorbent paper to remove residual liquid.
222. Wash 3 times in washing buffer. Tap the plate on absorbent paper to remove excess of PBS tween.
223. Add 100 μl of TMB solution (see "Reagent Setup").
224. Protect from light and incubate 15 min at room temperature.
225. Stop the reaction by adding 100 μl of 2N sulfuric acid.
226. Read the optical density with a plate reader at 450 nm.

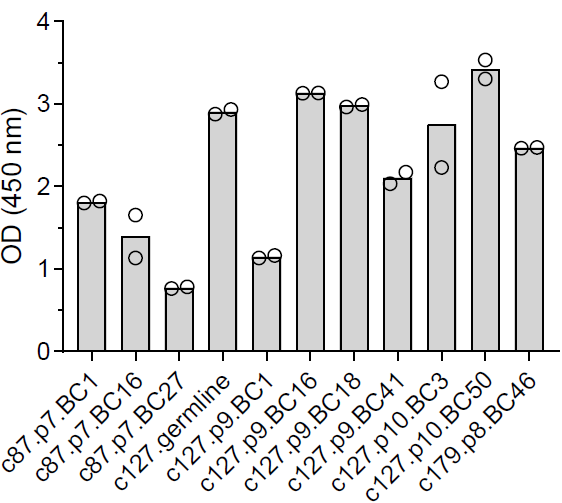

**ELISA control of production:** This ELISA allow us to estimate production of antibodies. In this graph. all supernatants of culture produced antibodies with some differences. This ELISA is performed to save money if some supernatants did not produce antibodies at all. Then. this ELISA allow to select only supernatants with antibodies for the purification with protein G beads because they are expensive.

Q. **Recovery of culture supernatant and preparation of 50% protein G beads - Timing 1h00**
227. After the ELISA results, collect the flasks that produced antibodies.
228. Transfer the contents of each flask into annotated 50 ml tubes.
229. Centrifuge the culture supernatants at 5000 g for 30 min at 4°C.
230. Filter the culture supernatants through 0.22-micron steritop filters using a vacuum pump system.
231. Transfer the filtered supernatant to a new 50 ml tube.

***Pause point:*** *Supernatant can be stored for few months at -20°C. For long-term storage, it is better to purify antibodies.*

232. Wash the protein G beads in PBS by centrifuging 5 min at 600 g. Repeat this step by changing the PBS.
233. Prepare the diluted protein G beads at 50%. The stock is around 70% of beads. Per tube you need 500 μl of 50% beads. (50% of protein G beads. 50% (vol/vol) of 1 × PBS).
234. Add 500 μl of diluted protein G beads into each tube.

**Tips:** Be careful to do this quickly as the protein G beads sediment to the bottom by gravity.

235. Close the tubes and add parafilm around the cap.
236. Put the tube on a rotator at 15 rpm at 4°C overnight.
R. **Antibodies purification – Timing 4h00**
237. Collect tube and centrifuge at 600 g for 5 min.
238. Aspirate the supernatant from tube using the vacuum pump system and leave about 1 ml of medium so as not to aspirate the beads.
239. Hydrate chromatography columns with 3 ml of PBS.
240. Resuspend beads with the remaining supernatant in the tube and put the supernatant with the beads on the columns.
241. Make two washes with about 3 ml of PBS.
242. For the second wash put a cap at the end of the column except for the first one to let it decant.
243. Once the first tube has decanted. remove the cap from the second tube and so on.
244. Elute each column with 1.2 ml of glycine buffer in four fractions of 300 µl in 1.5 ml tube containing each 30 μl of Tris buffer pH 8 (see "Reagent Setup").

**Tips:** The time that antibodies are at low pH should be minimized as much as possible.

245. Vortex briefly the tube to neutralize the pH.
246. Repeat steps 243 to 245 for the other columns.
247. Measure the concentration of the protein IgG by optical density with nanodrop at 280 nm and pool each fraction with detectable antibody.

*See troubleshooting*

S. **Antibodies concentration – Timing 1h00**
248. Hydrate 30KD amicon columns with 400 µl of PBS.
249. Load 400 µl of each amicon columns. Centrifuge at 14.000 g for 5 min. Repeat this step 4 time until no more antibodies are present in the Eppendorf tubes.
250. Wash columns twice with PBS and centrifuge at 14.000 g for 5 min.
251. Wash once columns with 0.04% sodium azide PBS.
252. Elute by inverting the column into a new tube and centrifuge at 3000 g for 2 min.
253. Measure the concentration of the protein IgG by optical density with nanodrop at 280 nm or equivalent and SDS-PAGE analysis in reduced and non-reduced condition.
254. This solution can be used immediately or stored at 4°C for month or at -80°C for years.
