## Supplementary File 3 for "FB5P-seq-mAbs: monoclonal antibody production from FB5P-seq libraries for integrative single-cell analysis of B cells"

**Troubleshooting table :**

| **Step** | **Problem** | **Possible reason** | **Solution** |
| --- | --- | --- | --- |
| 131 | The resulting libraries are contaminated by repetitive peaks of regularly sized DNA (“hedgehog” appearance of BioAnalyzer trace). | 5’-end TSO concatemers may form during RT. Final sequencing library quality will be negatively impacted by the presence TSO concatemers. | Perform a second round of library purification with a 0.7X ratio of CleanNGS beads. |
| 158 | No amplification of IGH or IGK chain by PCR | Multiplex PCR primers do not recognize the V region clonotype  Not enough preamplified cDNA in well  No amplification even after increasing the amount of cDNA or designing clone-specific primers | Design clone-specific primers targeting the exact sequence obtained from FB5P-seq  Increase the amount of preamplified cDNA (up to 8µl)  Have the IGHV and IGKV cDNA synthesized by a provider (*e.g.* Twist Bioscience or IDT). |
| 182 | No colony after SLIC reaction | Wrong insert:vector ratios | Check the concentration of insert and vector  Use classic restriction enzyme-based cloning instead of SLIC |
| 194-195 | PCR product from bacterial colonies with unexpected mutations | Mutations introduced by the PCR reaction | Test new colonies and run a new PCR |
| 247 | Low antibody yield | Antibody production may decrease with too many passages of the Expi293^TM^ cells.  Some antibodies are expressed at lower levels due to their structure or other unknown factor.  Low purity of plasmid DNA midi preparation | Thaw another batch of cells for transfection  Increase the number and culture volume of Expi293^TM^ cells  Repeat plasmid DNA midi preparation |
